# Uncovering persistent biases in human path integration by separating left and right trials

**DOI:** 10.1101/2025.09.24.678205

**Authors:** Jonas Scherer, Martin M. Müller, Anabel Kroehnert, Martin Egelhaaf, Olivier J. N. Bertrand, Norbert Boeddeker

## Abstract

Navigation ability strongly varies between humans. Careful analysis of errors occurring in navigation, or indeed any cognitive process, offers insight into the underlying mechanisms at different levels. While analyses at the individual level allow nuanced identification of error persistences across conditions and time, the population level facilitates generalisation but precludes conclusions about lower-level phenomena.

Regarding the critical navigation mechanism of path integration – the continuous tracking of navigated paths for self-localisation – previous studies have focused on populationlevel analyses, revealing systematic errors in estimating the travelled angles and distances. However, at the individual level, there are indications that people also possess left or right biases that are classically overlooked when pooling left and right trials. Therefore, we carefully investigate individual path integration errors in (1) a re-analysis of data from several influential human navigation studies, and (2) our own virtual reality path integration experiment. For both, we confirm time-persistent individual side biases, but find no evidence for consistent errors at the population level, suggesting that important aspects in human navigation performance might be overlooked by averaging across sides.

## 1 Introduction

Spatial navigation is a crucial cognitive function essential for many daily tasks, from finding one’s way in a city to navigating a room. However, sometimes we get lost. Errors made during navigation – alike those made in other cognitive tasks – offer valuable insights into the underlying processes. Analysing the structure of the errors enables specific affected cognitive mechanisms to be identified, including perception, memory, attention, or decision-making. Importantly, not all errors are random. Any errors that are persistent across different time points, conditions, sides, individuals, throughout populations, or any other experimental dimension can reveal stable characteristics of the cognitive mechanisms involved. Here, we analyse behavioural errors in spatial self-localisation along various experimental dimensions, and find evidence that population-level conceptions in the literature do not accurately capture persistent individual navigation biases.

Self-localisation is informed by multiple cognitive processes. In environments such as dark, dense spruce forests, most navigation methods, such as using landmarks, celestial objects, or background features, prove ineffective. However, we can continuously track navigated distances and angles over time which allows us to derive a sense of self-location by summing up motion signals from visual, vestibular, and proprioceptive sensory input; a process that is called path integration (PI) [1, 2, 3, 4].

In order to assess the performance of PI in human navigation, ‘homing’ tasks, in which participants must find their way back to a previously visited location, have received much attention in the spatial cognition literature [5, 6, 7, 8, 9, 10, 11]. One of the most prominent homing tasks is the triangle completion task [4, 6, 7, 12] in which a participant is guided away from a home location along an outbound path consisting of two sides of a triangle, and then asked to point to, or actively return to, the initial starting position, hence, completing the triangle (Fig. 1a). To successfully perform the triangle completion task, participants must first accurately estimate the rotation angle from the end of the outbound path to the starting point and then the physical distance between the two points. When scenarios without other spatial cues are provided, a triangle completion task isolates PI and allows its specific spatial errors to be quantified. Homing performance in such a task has been shown to depend on a variety of factors including triangle size, triangle shape, objects in the surround, and optic flow [3, 4, 6, 7, 12, 13, 14, 15].

**Figure 1.**
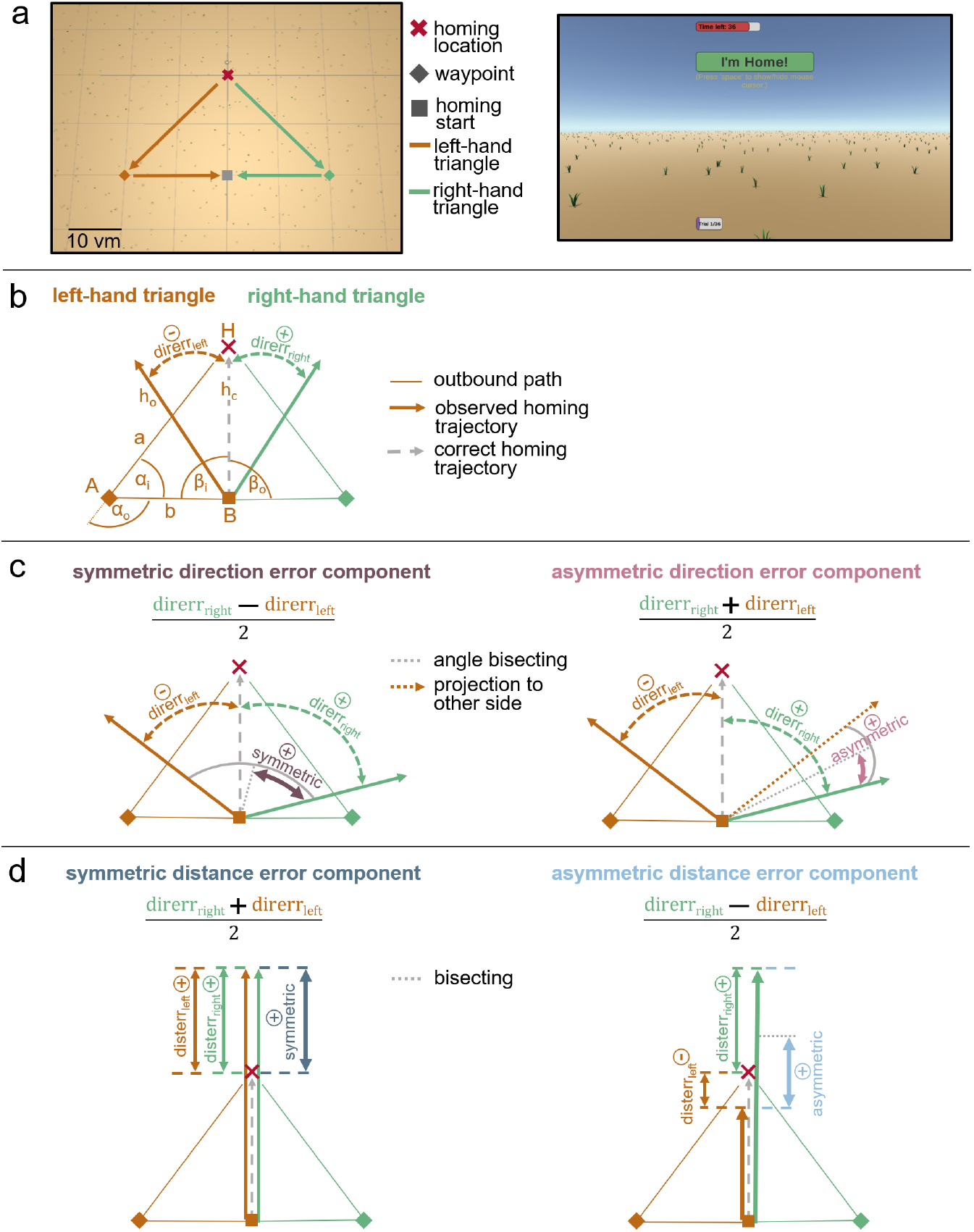
Experimental paradigm (a), definition of walked triangles (b), and analysed error components in direction domain (c), and distance domain (d). (a) Participants are tested in left (orange) and right-hand (green) triangle completion tasks in a desktop VR task (left: top view; right: screenshot of first-person view with red timeout bar, green response button, and purple progress bar at the bottom). Scale is given in virtual meters (*vm*), that correspond to actual meters in the VR. (b) Participants are guided along an outbound path on two legs, *a* and *b*, of a triangle with inner angles *α*_*i*_ and *β*_*i*_, and have to freely return to their initial home location (*H*). On their way they pass corner *A*, turn through angle *α*_*o*_ and reach corner *B* where they start free homing. To get to the correct homing trajectory *h*_*c*_ participants have to turn through angle *β*_*o*_. The observed homing trajectory *h*_*o*_ is not perfect, resulting in an observable direction error (*direrr*), defined as negative when deviating to the left (⊖ *direrr*_*left*_), or as positive when deviating to the right (⊕ *direrr*_*right*_) from the participant’s ego-perspective. Direction errors are calculated based on response endpoints. (c) Participants show symmetric (left, dark magenta) and asymmetric (right, light magenta) direction error components. The symmetric direction error component is the shared degree of over- or underturning in left and right-hand trials. The asymmetric direction error component describes the side tendency to the left or right side from ego-perspective of the participant. (d) Additionally, participants show symmetric (left, dark blue) and asymmetric (right, light blue) distance error components. Distance errors (*disterr*) are defined as negative when not walking far enough (*⊖ disterr*_*right*_ in right panel), or as positive when walking too far (e.g. *⊕ disterr*_*left*_ in right panel). The symmetric distance error component is the average degree of over- or undershooting the correct homing distance in left and right-hand trials. The asymmetric distance error component describes the tendency to walk further for one triangle side than for the other.

Beyond such task-level factors, navigation performance is also shaped by individual differences that emerge throughout the cognitive mechanisms involved in human navigation. These differences include individual preferences in directional strategies (survey versus route-based approaches) [16], and in reference frames (egocentric or allocentric perspective) [17, 18, 19]. Abilities to form mental representations of space vary substantially, both in real-world and virtual environments, with variability in detail and complexity [20, 21, 22]. Moreover, factors such as age, gender, upbringing environment, and geographic location play significant roles in shaping navigational abilities [23].

For triangle completion tasks some characteristic PI errors have been identified at the population and individual level. Firstly, population-wide errors in PI consist of two sub-components, an underestimation of walked distances, and of angles turned through [1, 3, 4, 12, 13, 24, 25, 26, 27, 28, 29, 30]. These misestimations of walked distances and turned-through angles make people typically overshoot small target angles and vice versa undershoot large target angles which can be observed in a regression to the mean for distance and angle responses in triangle completion tasks [1, 3]. Overshoots of target angles were also observed in spiders, ants, bees, hamsters and dogs (see [31] for a review), and were interpreted as a functionally useful behaviour in desert ants because they cause the return path to cross the first leg of the outbound journey, enhancing the chance to identify familiar landmarks [32]. However, such inaccuracies are typically only analysed at the population level, leaving open whether the reported angle and distance misestimations are truly systematic across individuals.

Secondly, there are participant-specific, individual side tendencies. Side tendencies have mainly been discarded in past research, either because left–right deviations were assumed to reflect random noise and thus examined only at the population level, where individual effects cancel out [1, 3, 4, 7, 11, 14, 33, 34], or because they were actively corrected for in the analysis [6]. This assumption may have obscured the possibility that participant-level biases contribute meaningfully to the error signal. In this regard, Jetzschke et al. investigated differences between left and right turns in freely walking, blindfolded participants without pooling at the population level, and first proposed the term ‘idiosyncratic error’ to describe individual tendencies to the left or right in their experiment [2]. They concluded idiosyncratic side biases to be persistent across time because participants estimated left and right turns differently by up to 57° (14.2 ± 14.3, mean±standard deviation (sd)) in an active and a passive turn estimation task that were conducted a few weeks apart [2]. Whether these idiosyncratic biases extend to PI performance in more complex triangle completion tasks and persist across different path shapes remains unknown. Previous work on desert ants suggests that they ‘turn as often to the right as to the left,’ which makes ‘an overall directional bias unlikely to develop’ [35]. Similarly, a study examining people’s ability to walk in a straight line over extended distances found that veering does not stem from a systematic directional bias but rather from random fluctuations in ‘the subjective sense of straight ahead’ [36]. Revealing persistent directional biases in humans performing triangle completion tasks would challenge these assumptions and provide new insights into how direction and distance estimates are calculated during path integration. Such findings would imply that PI is not a neutral, unbiased process but instead incorporates individual asymmetries in sensory signalling or higher-level processing, tracing some proportion of errors back to consistent distortions rather than random noise.

Finally, there are true random errors that are assumed to be caused by noise in the navigation system, which affects response precision (i.e. variance). They might be caused by noise originating in encoding sensory input [4, 25], in path calculations [37], or in motor execution [2, 3, 14, 38].

In summary, angular overshoots have been attributed to an adaptive behavioural function that facilitates the recognition of familiar landmarks. In contrast, undershoots or persistent directional left/right biases appear maladaptive and counterintuitive, yet their existence in human navigation remains unexplored.

In this study, we analyse individual misestimations of angles and distances in PI at various experimental dimensions. As individual PI side tendencies have often been ignored or treated as noise, we collected evidence for their existence in the available literature on human navigation, and re-analysed, with the consent of the respective authors, data from 11 influential, published studies that included some form of left and right hand trials [1, 2, 3, 4, 6, 7, 11, 33, 39, 40, 41]. The results reveal tremendous variance between individuals in all analysed studies.

In addition, we conducted dedicated experiments to analyse persistence of individual PI side tendencies across the experimental dimensions of time and triangle shape in our own virtual reality (VR) triangle completion task. For time persistence, we check correlations between performance in the same experimental setting in two sessions conducted at least three weeks apart, and for persistence across triangle shapes additionally with a triangle of a different shape. To resolve differences between the turning directions, participants are tested in triangles that contain either only left turns (left-hand triangle) or only right turns (right-hand triangle). If the previously observed idiosyncratic side tendencies [2] extend to triangle completion tasks, they would cause rotation errors to differ depending on the direction in which the triangle is traversed. We hypothesise to find a side-dependent error component, that hence is asymmetric between both triangle sides, and a side-independent error component, that is symmetric between both sides. From participants’ behaviour in an adequate number of repetitions in left-hand and right-hand triangles, we extract the *‘asymmetric error component’* that is dependent on rotation direction, and the side-independent *‘symmetric error component’*, by explicitly not pooling left and right trials. With this approach, we aim to determine to what extent individual biases contribute to the group-level errors reported in previous research.

The aim of this study is to answer the following questions: Are misestimations of angles really systematic and valid for all individuals? Is PI performance of individuals independent of side? Are individual side biases persistent across time and triangle shape? Our hypotheses are based on the study by Jetzschke et al. [2]. First, Jetzschke et al. demonstrated strong and highly variable directional biases in rotation estimation between participants. Since rotation estimation is a core component of PI, such biases are likely to manifest in PI performance as well. We therefore hypothesise that previously reported systematic PI errors at the population level are overstated, and that individual PI performance in triangle completion tasks depends on rotation direction. Second, the authors provided evidence that such biases persist across time and across different tasks, although their magnitude varied between tasks. Accordingly, we hypothesise that individual PI biases are persistent across time and across different triangle shapes.

Uncovering individual side biases in human PI behaviour that are persistent across conditions would be striking, as it could reveal a previously overlooked feature of spatial processing, with theoretical and empirical implications for future research on human navigation: Firstly, from a theoretical perspective, PI biases help us to better characterise individual navigation behaviour and provide insights that can guide future studies aimed at uncovering the underlying neural mechanisms. Secondly, from an empirical perspective, we pinpoint the size and type of biases in human PI in order to formulate best-practice advice on how to account for their effects in the design of future PI experiments. Our analysis at the individual level reveals individually persistent patterns in human PI and challenges established conceptions about population level errors.

## 2 Results

To analyse persistent errors in human PI at various experimental dimensions, we explore individual symmetric and asymmetric error components in two ways: first, we re-analyse data from influential, recent human navigation studies to assess generalisability of personal PI errors across tasks, and, second, we assess their persistence across time and path geometries in our own VR triangle completion task.

The symmetric and asymmetric individual error components are calculated based on signed direction errors from a VR triangle completion task (Fig. 1a and b). We define the signed direction error (*direrr*) based on the endpoint of a participant’s trajectory when returning to the initial home location. The signed direction error is the difference between the correct and the observed homing direction with a positive sign indicating right deviations, and a negative sign indicating left deviations, from a participant’s ego perspective. To resolve differences between turning directions, we separately consider trials with left turns, or right turns (Fig. 1b). The degree of the symmetric error component (*err*_*sym*_) is calculated by:

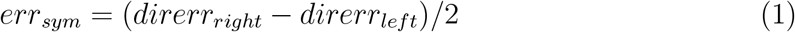

where _*right*_ and _*left*_ indicate triangle sides. This component is the shared degree of over- or underturning in left and right turns. Figuratively, one can illustrate the symmetric error component as half of the angle between the walked directions in a left-hand and a right-hand trial (Fig. 1c, in magenta).

The asymmetric error component is calculated by:

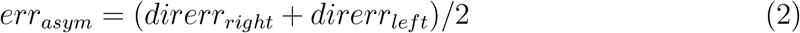

This component describes the tendency of a participant towards the left (negative sign) or right (positive sign) side. One can think of the asymmetric error component as the difference between the tendencies to over- or undershoot left or right turns. This can be illustrated as half of the angular difference between the vector pointing into the walked direction of a right-hand trial, and the projection of the vector of a left-hand trial to the other side (Fig. 1c, in light magenta).

In summary, the symmetric error component indicates the degree of over- or under-shooting the correct target angle on both sides, and the asymmetric error component reflects a directional bias to one side. Both error components are aggregated and averaged across trials (see section ‘Statistics’ in ‘Methods’ for details).

### 2.1 Re-analysis of data from published studies

We analysed datasets of 11 published human navigation studies that allow us to calculate symmetric and asymmetric error components from separate left and right trials [1, 2, 3, 4, 6, 7, 11, 33, 39, 40, 41]. Studies were selected based on a systematic literature search focused on citation counts, following the basic procedure reported in the PRISMA 2020 statement [42] (see supplement Fig. S1 for flow diagram). Datasets and consent for re-analyses were acquired from the respective authors via email. From directional errors, we calculated error component sizes strictly at the individual level and only eventually averaged across all conditions of a study. Detailed descriptions of each single study’s re-analysis are provided in the supplementary material (supplement S1). None but one [2] of the analysed studies was designed to explicitly investigate individual error components, but mostly focused on different experimental conditions instead of high individual repetition counts, meaning that the power of the single study re-analyses is low. Here, we focus on the collective evidence of the re-analysed studies for the existence of individual symmetric and asymmetric error components in human PI.

In all of the re-analysed studies we find a large variance in the symmetric and asymmetric error components, with 10*/*11 studies showing even larger inter-individual variance in the asymmetric error component than the present study (Fig. 2). Although 10 studies show a mean positive shift of the symmetric error component, reflecting average over-shooting of the correct homing angle, all of these studies also contain a considerable number (7% − 56%) of participants that undershoot the correct homing angle. The distribution of symmetric error components across all studies is significantly shifted to positive values with a mean of 10.6° (*N* = 275 participants in 12 studies, 95%-confidence interval (ci): [6.1, 14.9]). The distribution of asymmetric error components across all studies shows no significant difference from zero at the population level (*mean* = 1.4°, 95%-ci: [−1.2, 3.8]); however, the absence of a population shift from zero masks significantly larger individual biases in both directions that largely cancel each other out (Fig. 2 percentage bars).

**Figure 2.**
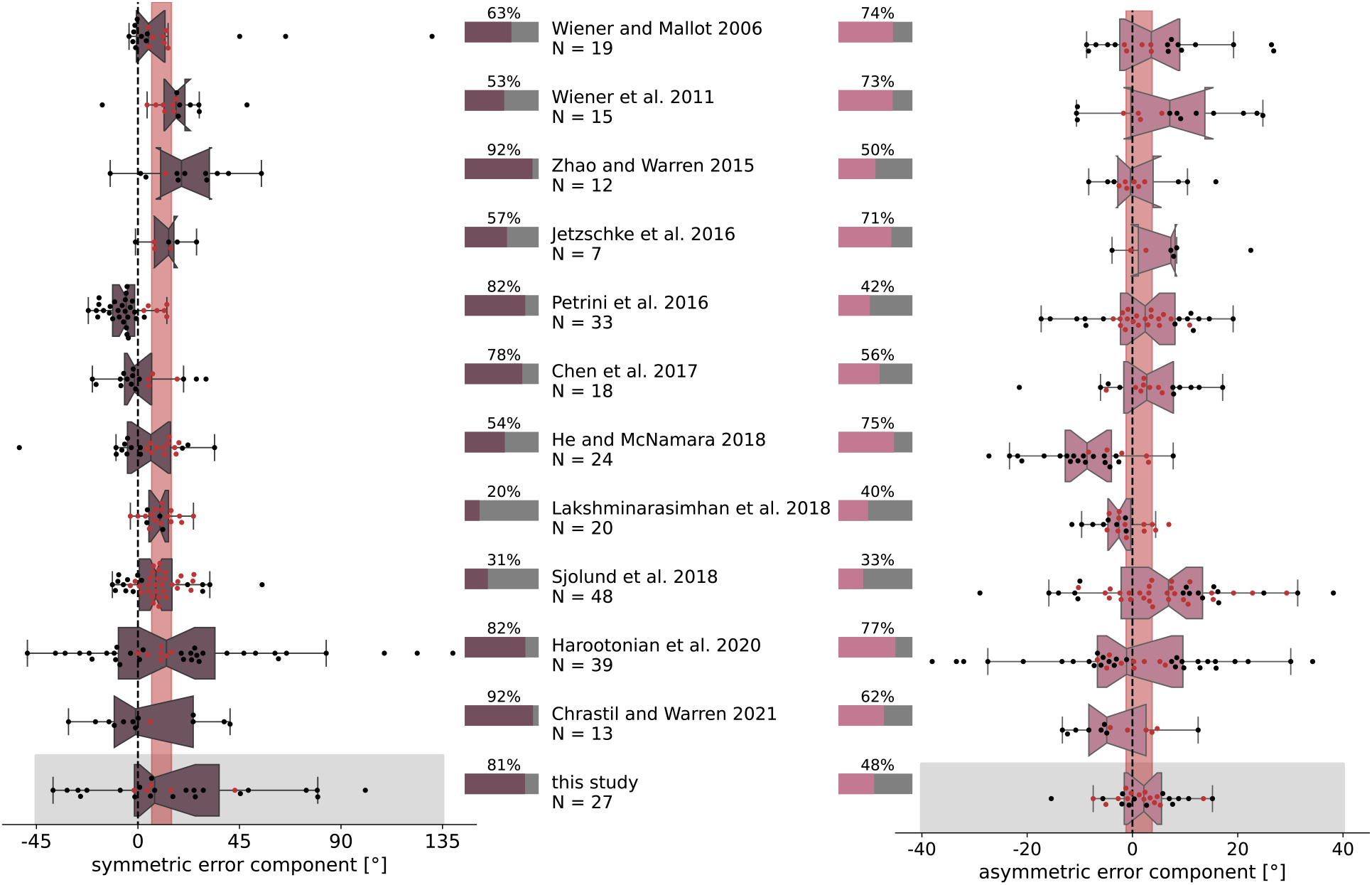
Symmetric and asymmetric error components vary strongly between individuals in 11 influential, published human navigation studies, and in this study (grey boxes). Symmetric (left, dark magenta) and asymmetric (right, light magenta) error components were calculated based on directional errors strictly at the individual level and only eventually pooled across all conditions in a given study (see supplement S1 for details). Boxplot notches indicate 95%-ci around the median for each study. Vertical red bars indicate 95%-ci around the mean across studies (symmetric: 10.6, [6.0, 15.0]; asymmetric: 1.4, [−1.2, 3.8]), and emphasise the large discrepancies many individuals have from the average measure. The total number of analysed individuals (*N*) and the percentage of individuals that significantly (bootstrapping hypothesis testing) differ from the population mean (horizontal bars) are indicated for each study. Each dot represents the averaged symmetric or asymmetric error component size of one individual and is coloured black when significantly different from the population mean, and else red. Published study datasets were selected, following the basic procedure reported in the PRISMA 2020 statement [42] (see supplement Fig. S1 for flow diagram).

Additionally, we compared studies in which participants actively walked ([2, 3, 4, 6, 7, 11, 33, 39, 40]), or used a joystick for control ([1, 41], and present study) for differences in symmetric or asymmetric error components. Neither the symmetric nor the asymmetric error component shows a significant difference in size between active walking or joystick control (permutation difference test: symmetric error component: *p* = 0.095, 95%-ci: [−7.18, 6.74]; asymmetric error component: *p* = 0.869, 95%-ci: [−3.12, 3.13]), but sd differs significantly between active walking or joystick control studies for both error components (F-test: symmetric error component: *sd*_*walking*_ = 22.5°, *sd*_*joystick*_ = 27.5°, *F* (65, 205) = 1.46, *p* = 0.047; asymmetric error component: *sd*_*walking*_ = 12.2°, *sd*_*joystick*_ = 7.8°, *F* (205, 65) = 2.43, *p <* 0.001).

The symmetric error component is considered to reflect a population-wide tendency to overshoot small angles and undershoot large angles, and hence has been labeled the ‘systematic’ PI error [2]. However, the results of our re-analysis of data from published studies reveal tremendous variance between individuals in the symmetric error component, exposing it at least to some degree as an individual error component. Additionally, the asymmetric error component proves to be highly relevant in understanding individual navigation behaviour as the re-analysis of data from published studies shows large inter-individual variance. Hence, we designed a dedicated study to analyse persistent symmetric and asymmetric individual direction and distance error components in PI.

### 2.2 Results from our virtual reality triangle completion task

We calculated individual symmetric and asymmetric PI error components from healthy participants (*N* = 27) completing *n* = 30 repetitions of randomly shuffled left and righthand desktop VR triangle completion tasks. In a triangle completion task, a participant is guided along two sides of a triangle and then asked to return to the initial home position. The task was conducted in a VR desert environment without defined landmarks or background features and with only sparse randomly arranged grass tufts on the ground to create optic flow for visual estimation of self motion (Fig. 1a). Before testing, participants conducted training trials that differed from test trials by the fact that they were guided from the homing start position to their initial home position by an arrow to initially calibrate their perception of rotations in the VR.

#### 2.2.1 Directional error components are significant and persistent across time

We dissect the total direction and the total distance error made by participants on average across trials into the symmetric and asymmetric error components. Although the asymmetric error component has not been analysed in most previous research, we find that it accounts for up to 90% (mean across participants: 21%) of the total direction error, and 92% (mean across participants: 14%) of the total distance error (Fig. 3). Both error components reveal noteworthy variance between individuals in their proportion and size (Fig. 3 and 5).

**Figure 3.**
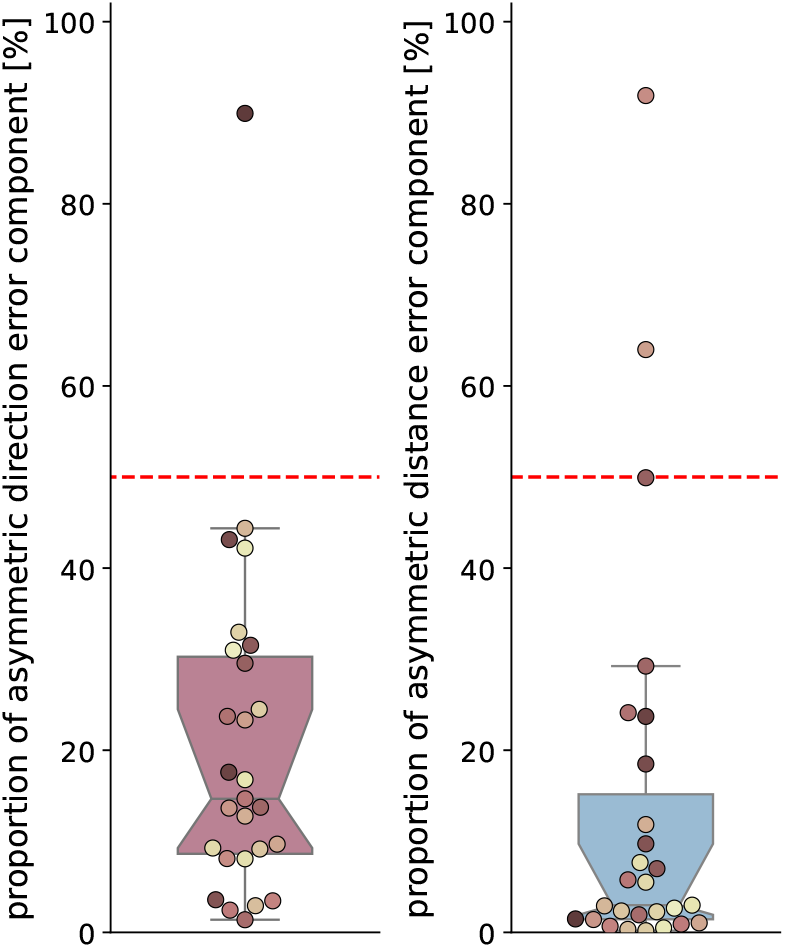
Proportion of asymmetric direction (left) and distance (right) error components in the total direction error obtained for all *N* = 27 participants in the main experiment of this study. The proportion of each error component in the total direction error is calculated by *err*_*asymmetric*_*/*(*err*_*asymmetric*_ +*err*_*symmetric*_). Boxplot boxes indicate quartiles, center lines indicate the median, notches indicate 95%-confidence intervals, and whiskers extend to the maximum and minimum within 1.5*IQR beyond the boxes. Dot colours are sorted from dark to light, corresponding from low to high asymmetric direction error component size in session one (see Fig. 5b), and are consistent across all figures. Shown is data of the empirical experiments of the current study. See Fig. 2 for an analysis of other published datasets.

We find that participants show deviations of different degrees between left and righthand trials, yielding observable symmetric and asymmetric error components (see Fig. 4 for an example participant). Some participants show a strong positive symmetric error component, meaning they overturn the correct homing angle for both triangle sides, while others underturn it. Some participants show a strong positive asymmetric error component, meaning they underturn the correct homing angle in left trials more than in right trials, while others behave vice versa. Strong error components are consistently visible across experiment days. All three effects are shown for an example participant in Fig. 4.

**Figure 4.**
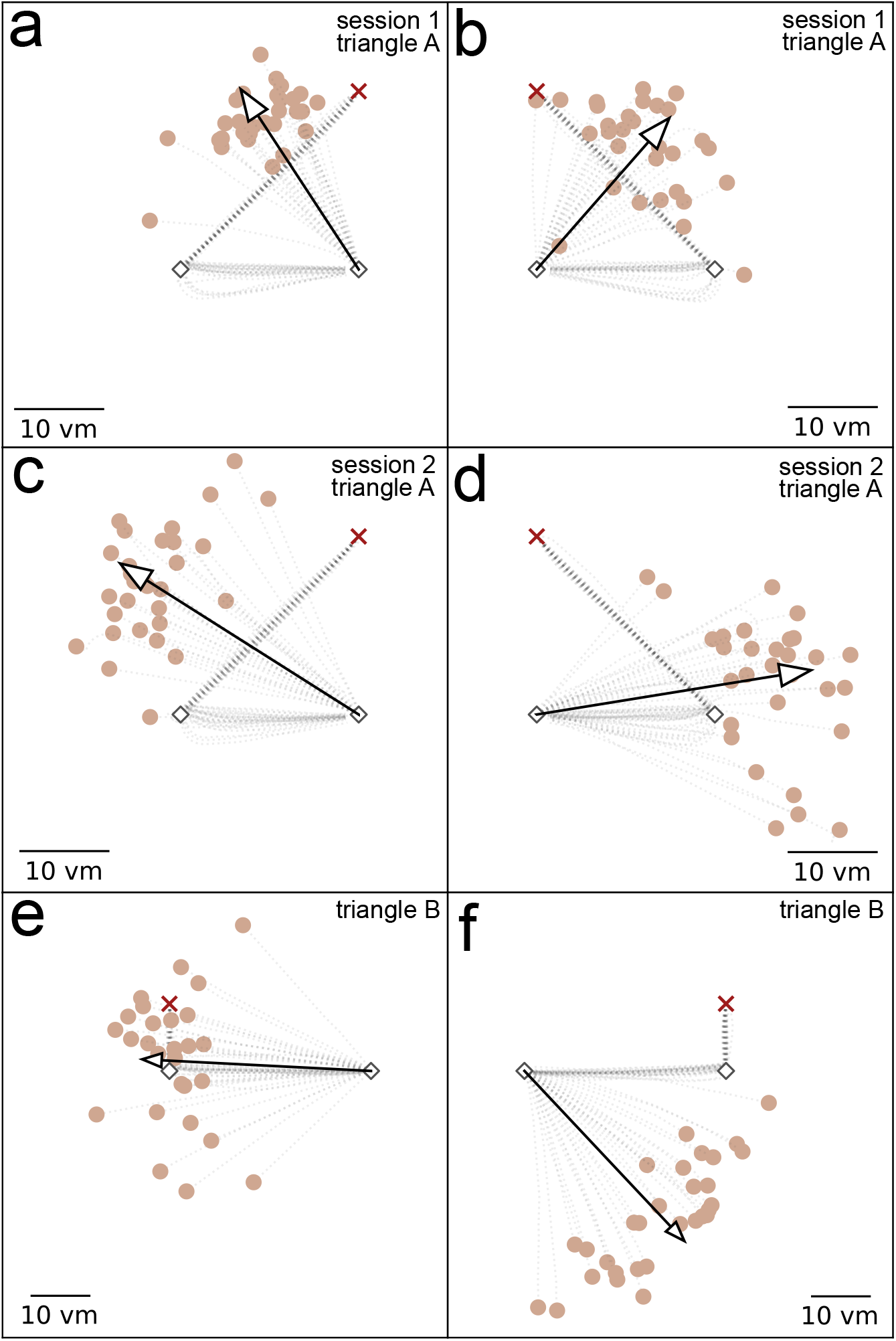
Homing performance of an example participants depends on triangle side and shape. The example participant overturns the correct homing angle in all conditions. Asymmetries between left and right-hand trials are apparent. The left column (a, c, e) shows left-hand trials, and the right column (b, d, f) shows right-hand trials. The upper row (a, b) shows trials in triangle A from session 1, the middle row (c, d) shows trials in triangle A from session 2 conducted at least 3 weeks later, and the bottom row (e, f) shows trials in triangle B conducted in between the other two sessions. Trajectory endpoints of single trials are depicted by dots. The depicted participant is uniquely identifiable across all figures in this paper as dot colour is consistent. Outbound and homing trajectories are indicated by light grey dotted lines. Red crosses indicate the home location, and black diamonds indicate waypoints. Arrows depict mean direction and distance. Scale is given in virtual meters (*vm*), that correspond to actual meters in the VR. Shown is data of the empirical experiments of the current study. See Fig. 2 for an analysis of other published datasets.

In the next step we calculated the symmetric and asymmetric error components for each individual participant for each of two sessions conducted at least three weeks apart (min= 21, mean= 35.7 days later) to analyse error component persistence across time (see section ‘Error component calculations’ in ‘Methods’ for details). We find participants with different degrees of negative (left) and positive (right) asymmetric error components, with the population distribution being centered around zero (Fig. 5b). Asymmetric error component sizes of session one and session two appear similar. The distribution mean of differences between the days is not significantly different from zero (bootstrapping hypothesis testing: *N* = 27, *p* = 0.414, 95%-ci: [−3.61, 1.1], *d* = −0.156). Importantly, our power analysis indicates that the study design was sufficient to detect differences of 2° or greater with 85% power, suggesting that any undetected difference, if present, is likely smaller than this threshold (see section’Power analysis’ for details). The test-retest reliability of the asymmetric error component was assessed using the intraclass correlation coefficient (ICC) with a two-way random effects model (ICC2). The ICC was 0.581 (95%-ci: [0.27, 0.78]), indicating moderate reliability (*F* (26, 26) = 3.73, *p <* 0.001).

**Figure 5.**
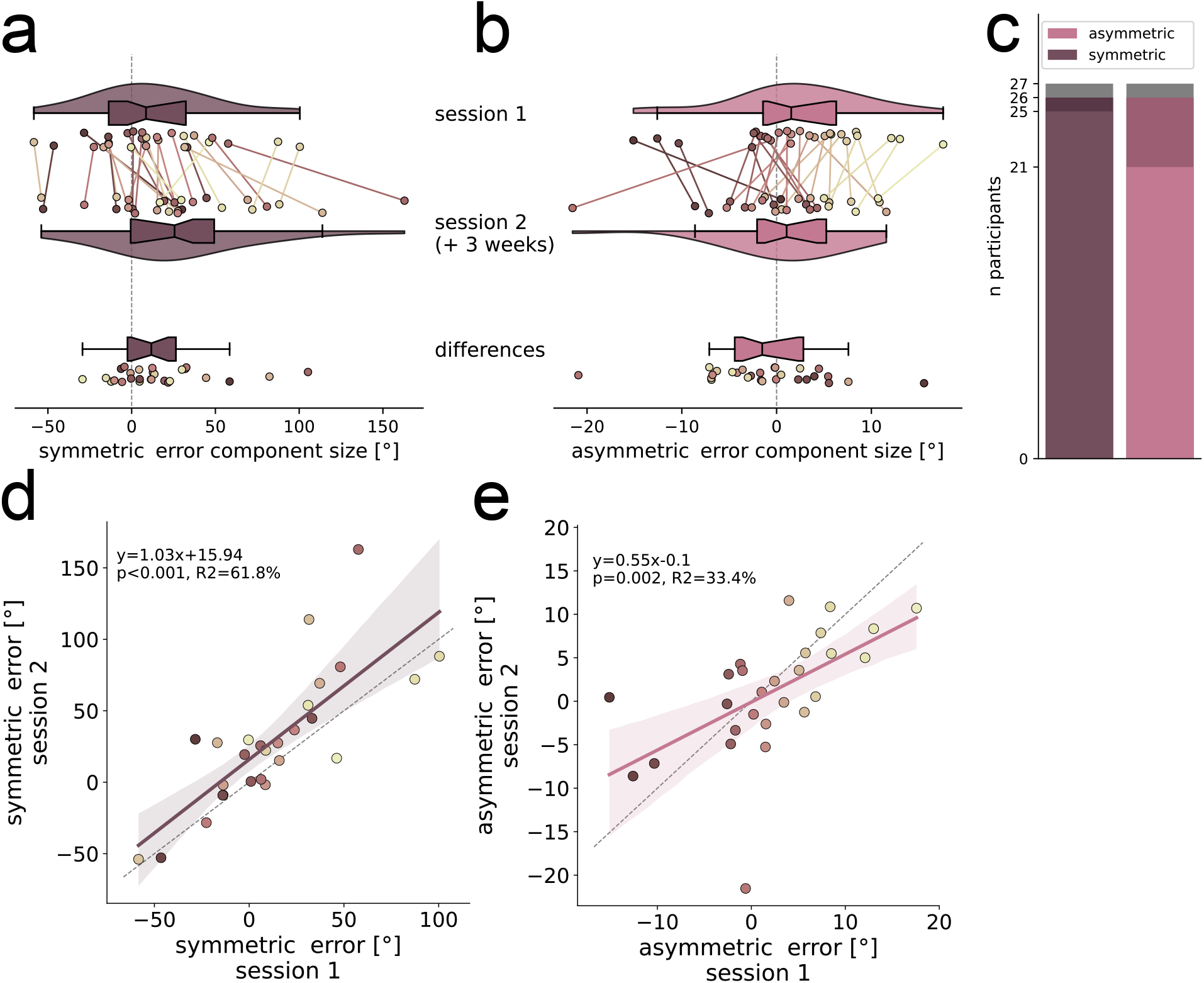
Symmetric and asymmetric direction error component sizes in session one and session two. (a) Symmetric and (b) asymmetric error component sizes, indicated by °, are calculated per individual participant (*N* = 27) for each of two sessions that were conducted at least three weeks apart and are indicated by different coloured dots. Lines of the same colour connect data points of individuals across sessions and indicate the amount of change. Dot colours are sorted from dark to light, corresponding from low to high asymmetric direction error component size in session one (see Fig. 5b), and are consistent across all figures. Violins depict kernel density estimations and contain boxplots with the boxes indicating quartiles, center lines indicating the median, notches indicating 95%-confidence intervals, and whiskers extending to the maximum and minimum within 1.5*IQR beyond the boxes. Additionally, the magnitude of differences between sessions is indicated. (c) Number of participants with significant (bootstrapped 95%-confidence intervals around mean) error components in one (dark colours) or both (light colours) sessions (see section ‘Error component calculations’ in ‘Methods’ for details). 26/27 participants show significant error components in at least one of two sessions. (d) Symmetric and (e) asymmetric error components are persistent across sessions. Linear regressions with 95%-confidence intervals are plotted in dark magenta (symmetric) or light magenta (asymmetric) and described by the line equation, coefficient of determination *r*^2^, and Pearson correlation *p*-value. Grey dashed lines show line with slope 1. Shown is data of the empirical experiments of the current study. See Fig. 2 for an analysis of other published datasets.

For the symmetric error component we find participants who systematically overturn the correct homing angle (positive), and also participants who systematically underturn it (negative) (Fig. 5a). This finding contradicts earlier studies that treat the misestimation of rotations as a population-wide effect [1, 3, 4, 12, 13, 24, 25, 26, 27, 28, 29, 30]. Surprisingly, the symmetric error component shows even larger variance between participants than the asymmetric error component. Symmetric error component sizes of session one and session two are similar with an average increase of 16.3° between the sessions, and the distribution of differences being significantly different from zero (bootstrapping hypothesis testing: *N* = 27, *p* = 0.004, 95%-ci: [7.1, 29.46], *d* = 0.562). The ICC for the symmetric error component was 0.712 (95%-ci [0.4, 0.87]), indicating good reliability (*F* (26, 26) = 7.28, *p <* 0.001). The variance between participants is larger in the symmetric than in the asymmetric error component (*N* = 27; sd: asymmetric: 7°, symmetric: 40.5°; *F* = 0.03, *df*_1_ = 26, *df*_2_ = 26, *p <* 0.001). Regression analyses between error component sizes of session one and session two reveal strong correlations for the symmetric as well as for the asymmetric error component (Fig. 5d and e). The error size in session one explains 62% and 33% of variance in error size of session two, for the symmetric and asymmetric error component, respectively. This implies that the symmetric and the asymmetric error components are individually persistent for sessions that lie at least three weeks apart.

Moreover, we use the *n* = 30 repetitions per triangle side to calculate a distribution of all 900 possible combinations of one left and one right trial for each of the two error components for each participant (see section ‘Error component calculations’ for details). Bootstrapping hypothesis tests on the error component distributions reveal that 26 of 27 participants show symmetric and asymmetric PI error components that are significantly different from zero in at least one session. Moreover, 78% of participants exhibit a significant symmetric error component, and 93% exhibit a significant asymmetric error component in both sessions (Fig. 5c). Thus, both error components can reliably be shown in most of the participants.

To validate our approach of calculating error components based on response trajectory endpoints of each trial we conducted the same analyses with only the initial heading direction of participants during the first 2 virtual meters (*vm*) of each homing path. The results of this analysis qualitatively resemble those displayed here (see supplement Fig. S2 for details).

Additionally, we analysed correlations between the error components and individual differences assessed via post-task questionnaires, including prior experience with computer games and handedness [43]. The symmetric error component significantly correlates with subjective gaming experience (*N* = 27, *y* = −7.6*x* + 62.6, *r*^2^ = 0.26, *p* = 0.006) and handedness (*N* = 27, *y* = 0.3*x* + 3.6, *r*^2^ = 0.16, *p* = 0.039), with right-handed participants tending to overturn and left-handed participants to underturn (supplement Fig. S3). Handedness findings should be interpreted cautiously given our limited sample of left-handed participants (3*/*27). No significant correlations were found for the asymmetric error component (supplement Fig. S3).

We find that both, the symmetric and the asymmetric error components make up a notable share of overall direction error in PI (Fig. 3) while being highly variable between participants and being persistent across two sessions conducted at least three weeks apart (Fig. 5).

#### 2.2.2 Directional error components are persistent across path shape

We tested most participants (*N* = 19) some days later (min= 5, mean= 21.7) also with a different triangle shape characterised by a larger homing angle (triangle A: 90°, triangle B: 162°; see Fig. 4). Asymmetric error component sizes were found to be similar in triangle A and triangle B (Fig. 6b), with the distribution mean of differences between the triangles not being significantly different from zero (bootstrapping hypothesis testing: *N* = 19, *p* = 0.449, 95%-ci: [−2.37, 6.04], *d* = 0.18). For the symmetric error component in triangle B we find more participants than in triangle A who underturn the correct homing angle (Fig. 6a). Symmetric error component sizes of triangle B are lower than for triangle A with an average decrease of 13.24° between the triangles, but the distribution of differences not being significantly different from zero (bootstrapping hypothesis testing: *N* = 19, *p* = 0.086, 95%-ci: [−27.14, 3.35], *d* = −0.39). Both error component sizes are similar across triangles A and B, while there is a tendency for the symmetric error component to decrease as the homing angle increases. This outcome is not surprising, as a regression to the mean effect reported in previous literature suggests that participants would underturn large angles. Although participants in our study mostly show lower symmetric error components with the larger homing angle, they still tend to overturn it.

**Figure 6.**
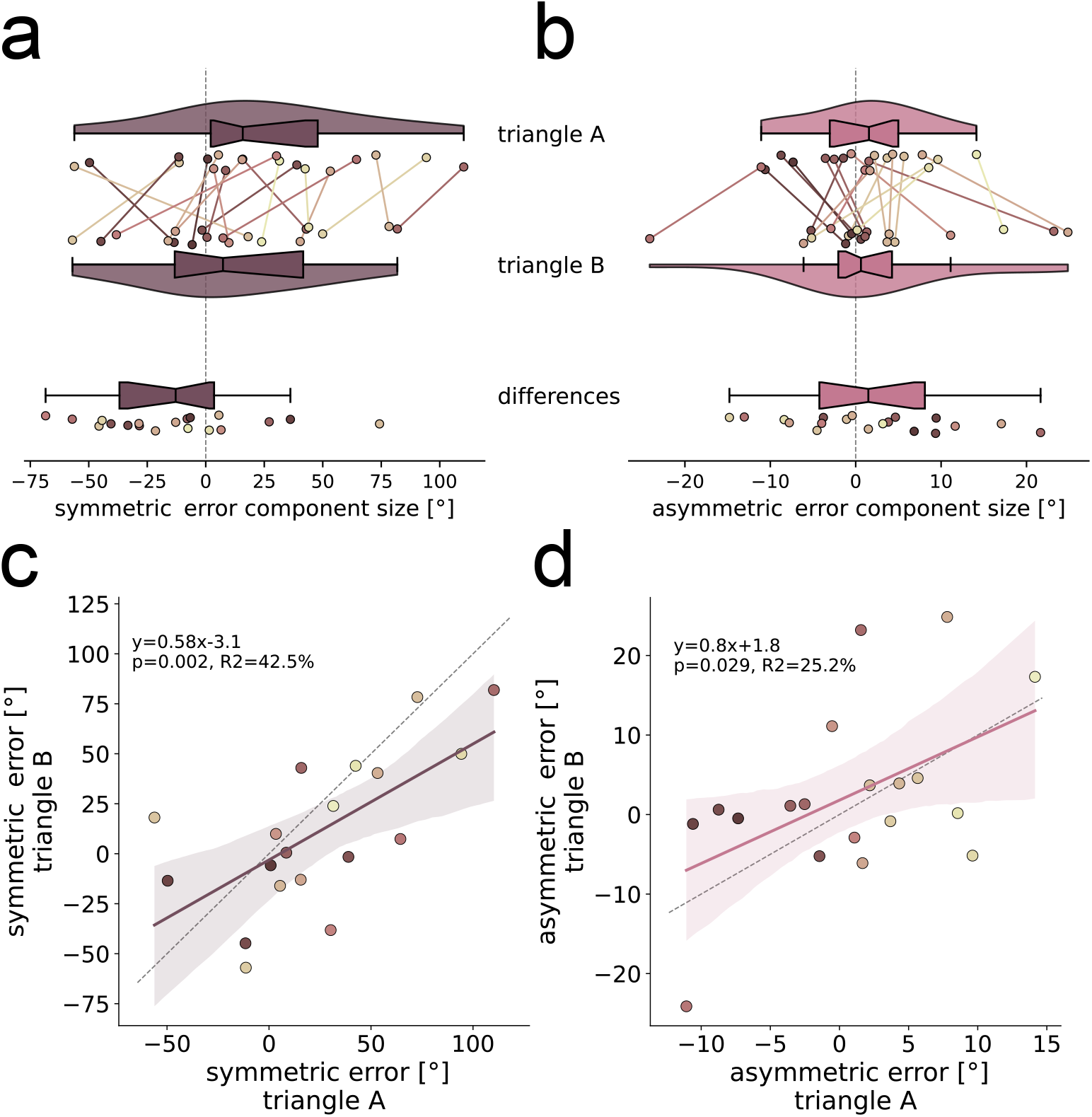
Symmetric and asymmetric direction error component sizes in triangle A and triangle B. (a) Symmetric and (b) asymmetric error component sizes, indicated by °, are calculated per individual participant (*N* = 19) for each of two triangle geometries and are indicated by different coloured dots. Violins that indicate data from triangle A show mean values across both experimental sessions conducted at least three weeks apart. Data for triangle B was collected in only one session. Only participants who were tested in both triangles are shown. Lines of the same colour connect data points of individuals across sessions and indicate the amount of change. Dot colours are sorted from dark to light, corresponding from low to high asymmetric direction error component size in session one (see Fig. 5b), and are consistent across all figures. Violins depict kernel density estimations and contain boxplots with the boxes indicating quartiles, center lines indicating the median, notches indicating 95%-confidence intervals, and whiskers extending to the maximum and minimum within 1.5*IQR beyond the boxes. Additionally, the magnitude of differences between sessions is indicated. (c) Symmetric and (d) asymmetric error components are persistent across triangle geometries. Linear regressions with 95%-confidence intervals are plotted in dark magenta (symmetric) or light magenta (asymmetric) and described by the line equation, coefficient of determination *r*^2^, and Pearson correlation *p*-value. Grey dashed lines show line with slope 1. Shown is data of the empirical experiments of the current study. See Fig. 2 for an analysis of other published datasets.

This regression to the mean effect is also evident in regression analyses between error component sizes of triangle A and triangle B which reveal significant correlations for the symmetric and the asymmetric error component (Fig. 6c and d). The regression slope for the symmetric error component is 0.58, meaning that the tendency to overturn for a given individual is lower in triangle B. The error size in triangle A explains 43% of variance in error size of triangle B, for the symmetric error component, and 25% for the asymmetric error component.

The symmetric and the asymmetric error components are individually persistent, but not equal, for two triangle shapes with strongly deviating correct homing angles of 90° and 162°.

#### 2.2.3 Distance error components are persistent

Besides error components in direction error, we can also calculate symmetric and asymmetric error components of distance estimation. The distance error is the deviation from the correct homing distance and defined as positive when participants walk too far and as negative when they walk too short. Hence, the symmetric distance error component describes the tendency of a participant to over- or undershoot the correct homing distance symmetrically for left and right-hand triangles (Fig. 1d). The asymmetric distance error component describes the tendency of a participant to over- or undershoot the correct homing distance for one triangle side more than for the other. A positive asymmetric distance error means the person walked further in right-hand triangles, while a negative value means they walked further in left-hand triangles.

We find participants with a positive and participants with a negative symmetric distance error component, meaning that some participants walk too far and some too short (Fig. 7a). The symmetric distance error component is persistent across the two sessions, indicated by a significant (*p <* 0.001) correlation with 50% of explained variance (Fig. 7d), with the error component size increasing in session two by 2.62 *vm* on average (bootstrapping hypothesis test: *N* = 27, *p* = 0.019, 95%-ci: [0.66, 5.13], *d* = 0.45; Fig. 7a). The ICC for the symmetric distance error component was 0.662 (95%-ci [0.37, 0.83]), indicating moderate reliability (*F* (26, 26) = 5.55, *p <* 0.001). We find variance between participants with some possessing a positive, some a negative asymmetric distance error component, but error component sizes are small (Fig. 7b). The asymmetric distance error component reveals no time persistence across the two sessions (*ICC*2: 0.183, 95%.ci [−0.22, 0.53], *F* (26, 26) = 1.43, *p* = 0.183) (Fig. 7e). There is no significant correlation of asymmetric distance error components across the two sessions (linear regression: *N* = 27, *r*^2^ = 0.03, *p* = 0.364).

**Figure 7.**
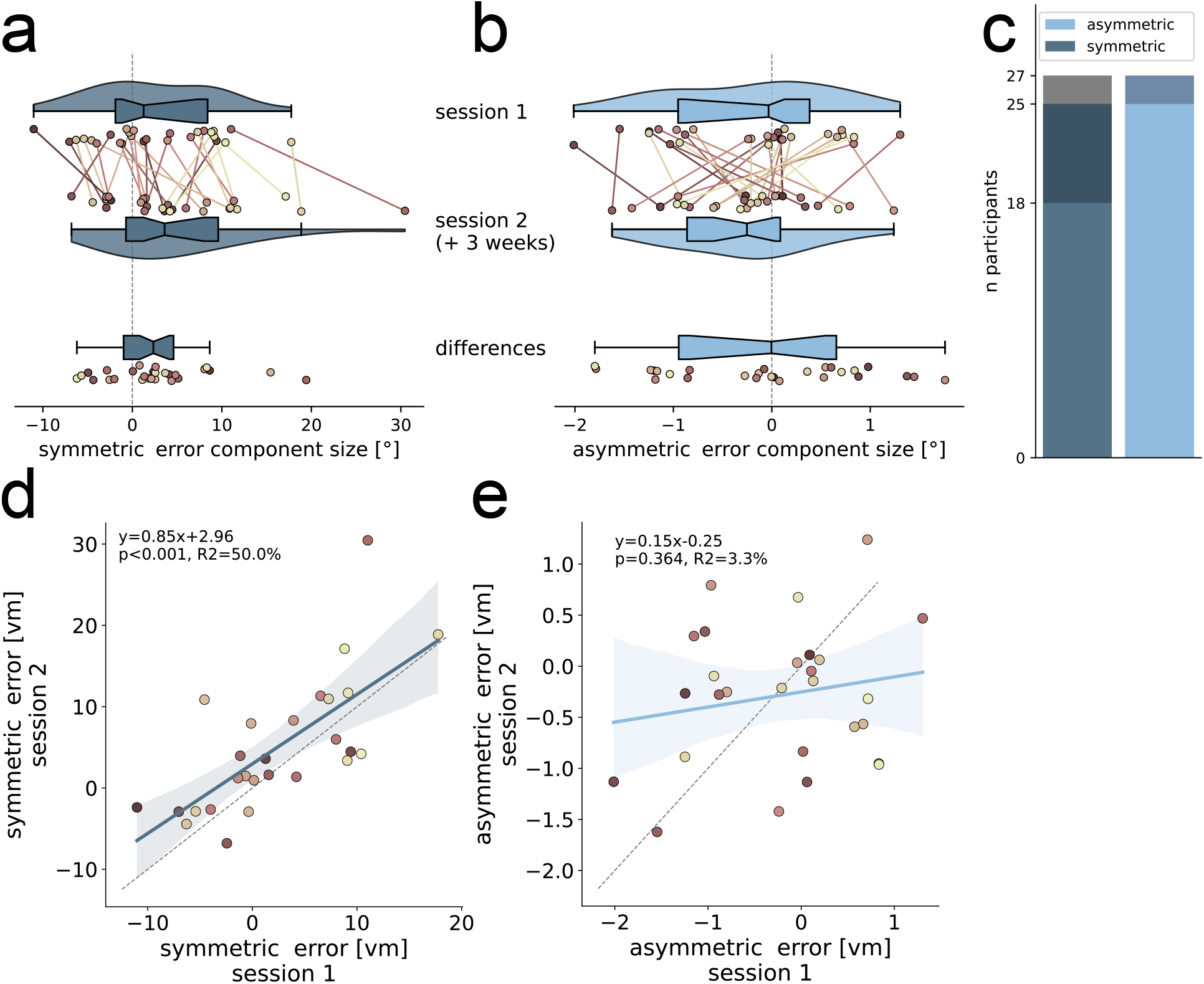
Symmetric and asymmetric distance error component sizes in session one and session two. (a) Symmetric and (b) asymmetric error component sizes, indicated by *vm*, are calculated per individual participant (*N* = 27) for each of two sessions that were conducted at least three weeks apart and are indicated by different coloured dots. Lines of the same colour connect data points of individuals across sessions and indicate the amount of change. Dot colours are sorted from dark to light, corresponding from low to high asymmetric direction error component size in session one (see Fig. 5b), and are consistent across all figures. Violins depict kernel density estimations and contain boxplots with the boxes indicating quartiles, center lines indicating the median, notches indicating 95%-confidence intervals, and whiskers extending to the maximum and minimum within 1.5*IQR beyond the boxes. Additionally, the magnitude of differences between sessions is indicated. (c) Number of participants with significant (bootstrapped 95%-confidence intervals around mean) error components in both (light colours), one (dark colours), or none (grey) sessions (see section ‘4.5.1’ in ‘Methods’ for details). (d) Symmetric error component is persistent across sessions, while (e) asymmetric is not. Linear regressions with 95%-confidence intervals are plotted in dark blue (symmetric) or light blue (asymmetric) and described by the line equation, coefficient of determination *r*^2^, and Pearson correlation *p*-value. Grey dashed lines show line with slope 1. Shown is data of the empirical experiments of the current study. See Fig. 2 for an analysis of other published datasets.

When comparing distance error components between triangle A and B, we find weak correlations across triangles (linear regression: *N* = 19; symmetric: *r*^2^ = 0.237, *p* = 0.035, asymmetric: *r*^2^ = 0.237, *p* = 0.034; supplement Fig. S4).

While both distance error components are highly variable between individuals, only the symmetric distance error component shows to be persistent across two sessions conducted at least three weeks apart.

#### 2.2.4 Symmetric direction and distance error components correlate

As our triangle completion task is designed in Euclidean space where directions and distances are spatially coupled, we correlate error components of direction and distance. We find a significant correlation of the symmetric direction error component with its equivalent for distance (linear regression: *N* = 27, *r*^2^ = 0.42, *p <* 0.001)(Fig. 8). For the asymmetric error components we find no significant correlation, but a trend, between direction and distance (linear regression: *N* = 27, *r*^2^ = 0.12, *p* = 0.073). Note that we also find a correlation between the measured direction and distance errors (no symmetric or asymmetric calculations included) for left-hand trials (*N* = 27, *r*^2^ = 0.29, *p* = 0.004), right-hand trials (*N* = 27, *r*^2^ = 0.35, *p* = 0.001), and both sides pooled (*N* = 27, *r*^2^ = 0.33, *p* = 0.002)(see supplement Fig. S5 for details).

**Figure 8.**
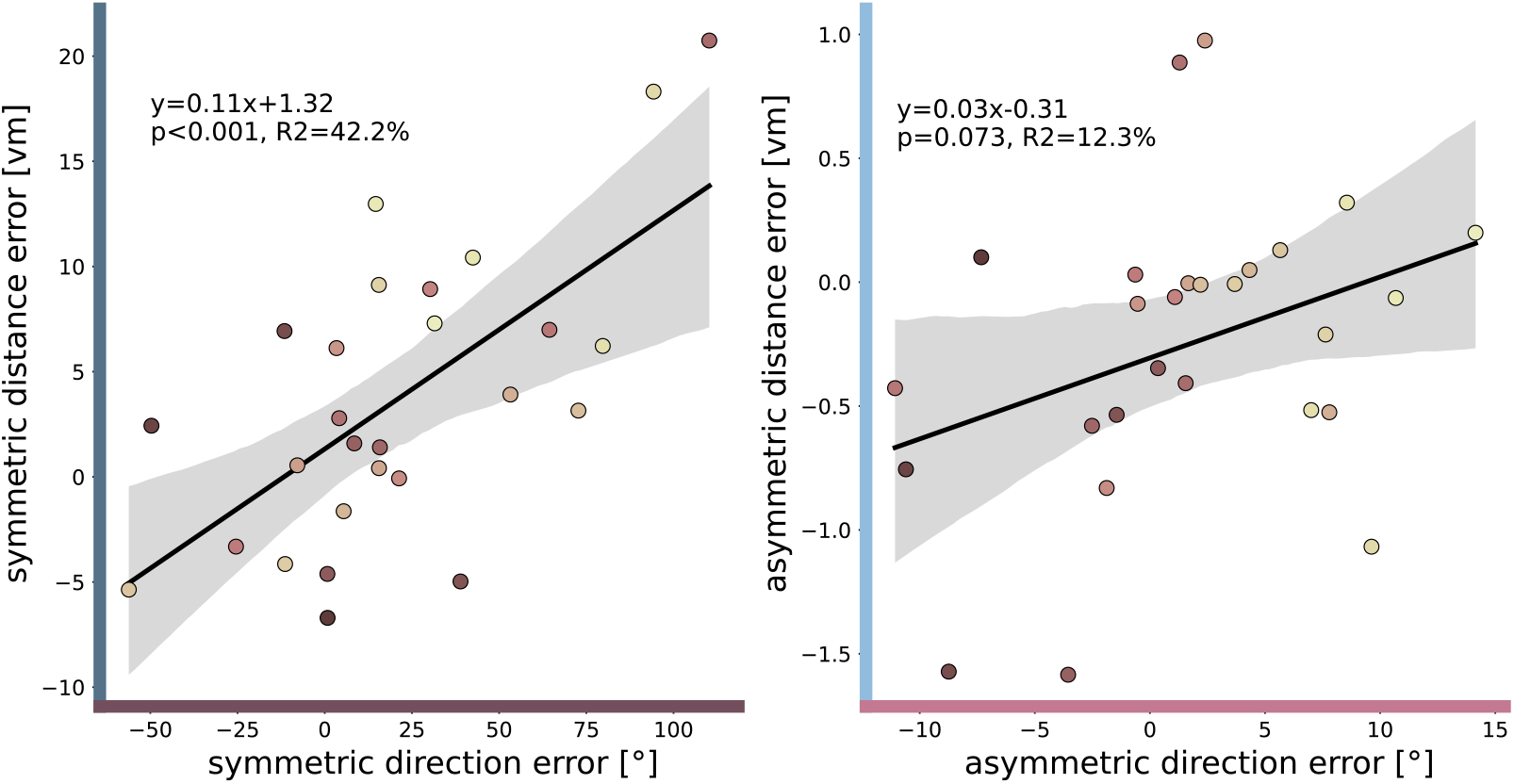
Symmetric and asymmetric error components for direction error and distance error are correlated. Error component sizes, indicated by ° for direction errors and in *vm* for distance errors, for individual participants (*N* = 27) are depicted by dots. Dot colours are sorted from dark to light, corresponding from low to high asymmetric direction error component size in session one (see Fig. 5b), and are consistent across all figures. Linear regressions with 95%-confidence intervals are described by the line equation, coefficient of determination *r*^2^, and Pearson correlation *p*-value. Grey dashed lines show line with slope 1. Shown is data of the empirical experiments of the current study. See Fig. 2 for an analysis of other published datasets.

## 3 Discussion

We investigated error components of path integration at various experimental dimensions to understand the role of population-, individual-, path-, time-, and side-dependent error components in human navigation. Building on that, we analysed the *‘asymmetric error component’* that is dependent on rotation direction, and the side-independent *‘symmetric error component’* with two approaches: (1) We re-analysed data of several recent human navigation studies for symmetric and asymmetric errors in turning behaviour. We find that participants from these studies show strong inter-individual variation in their error component sizes (Fig. 2). Across all re-analysed studies, the distribution of symmetric error components is significantly shifted positively by ≈ 10°, indicating a tendency of overshooting target angles across experimental designs. While individual differences account for most of the symmetric error component, the observed shift suggests that some portion is also attributed to systematic directional misestimation in PI that is consistent across many but not all of the participants. (2) We tested participants in an adequate number of triangle completion tasks in left-hand and right-hand triangles to investigate consistency of symmetric and asymmetric error components across time and triangle shapes. We find both error components to be subject to large inter-individual variance (Fig. 5a and b) and to be of significant size (Fig. 3). Participants in our own experiment persistently misturn the correct homing angles although they are initially trained to perform the task ideally by an arrow that fully guided them through the training triangle (see section ‘Experimental procedure’ in ‘Methods’). Persistent misturning is even more surprising when considering the random shuffle between left and right trials that we used to account for possible sequence effects of past trials on PI responses as shown by Harootonian et al. in their VR triangle completion task, which in our experiment might even have reduced asymmetric error components [4]. Thus, persistence in misturning indicates intrinsic mechanisms that cannot simply be calibrated by task training or randomly shuffled trials.

### 3.1 Watch out for ‘systematic’ path integration errors

Many previous publications have observed human PI behaviour at the population level and found that people ‘systematically’ overshoot small angles or distances, and undershoot large angles or distances (e.g. [1, 3, 4, 12, 13, 24, 25, 26, 27, 28]). Our re-analysis and our own experiment revealed no population-level consistency in either symmetric or asymmetric error components. The collective evidence from the present study, and the re-analysis of published studies, illustrates that behavioural measures at the population level do not necessarily reflect behaviour accurately at the individual level, which might lead to imprecise conclusions about underlying cognitive processes.

### 3.2 Individual differences need to be incorporated in path integration research and navigation models

We find no evidence for the symmetric or the asymmetric error component to be consistent at the population level. Neither the symmetric nor the asymmetric error component should be discarded when trying to understand individual human PI behaviour because both error components make up a notable proportion of the total observed error in direction and distance (Fig. 3). Both error components show strong correlations at the individual level in two sessions conducted at least three weeks apart. This means that symmetric and asymmetric error components in our experiment are time persistent (Fig. 5d and e). Additionally, the two error components are persistent across path geometries, although not being equal therein (Fig. 6c and d). Unlike previous research, which proposed that veering from walking straight was due to random fluctuations [36], we find evidence of persistent directional biases that may also affect straight walking consistently.

The symmetric error component size decreases for most participants for a triangle shape with larger correct homing angle (162° vs 90°). This size reduction is unsurprising as other triangle completion studies also found participants to underturn large angles and overturn small angles [1, 3, 4, 12, 13, 24, 25, 26, 27, 28]. Additionally, Wiener and Mallot pointed out that large angles above 140° in their desktop VR experiment seemed to be hard to perceive and integrate correctly [1]. Although most participants in our experiment overturn the correct homing angle for both triangle shapes, they do less so when having to turn through the larger angle (Fig. 6a). Since we observe no consistent overshoot of the target angle, our findings do not support the hypothesis proposed for ants that angular overshoots serve an adaptive function by facilitating intersection with the outbound path and enabling landmark recognition [32].

Individual asymmetric error sizes are not equal between triangle shapes but we find a significant correlation between asymmetric error sizes of both tested triangle shapes (Fig. 6d). Dedicated study designs with more triangle shapes are necessary to quantify how the asymmetric PI error component generalises and scales with homing angle and paths with more, or naturally curvy, segments. Quantifying such generalised symmetric and asymmetric error component terms also enables existing PI models, such as the ‘encoding-error model’ [3, 4, 25], or the more recent ‘generative linear-angular model of PI’ [14], or ‘dynamic Bayesian actor model’ [38] to be refined. Refining PI models in this respect might also reveal why the existing encoding-error model performs poorly for paths with more than three segments [3, 25].

### 3.3 Origins and implications of path integration biases

This study is, to the best of our knowledge, the first to systematically analyse PI asymmetries in a VR experiment. Past studies show clear evidence that optic flow information without salient landmarks or vestibular cues is sufficient for participants to perform basic PI tasks in good agreement with free-walking experiments [1, 13, 44]. The results of the present study also agree with several published free-walking experiments included in our re-analysis of data from published studies (Fig. 2). As Jetzschke et al. did in realworld free-walking, we find evidence for the existence of individual side tendencies also in VR [2]. This methodological transfer might indicate a shared fundamental mechanism influencing and biasing navigation in both realms, which can be investigated in future studies by comparing individual performance in desktop and active-walking VR. Namely, Jetzschke et al. blindfolded their participants and provided them with only body-based and no visual stimuli, and hence they suggest asymmetries to arise in the vestibular system. Conversely, in our study, we test participants in a desktop VR with computer keyboard control and provide only visual input and no body-based information, hence suggesting underlying asymmetries in the visual system. Since we validate behavioural asymmetries from real-world experiments in VR, this might also suggest an overarching biased process beyond perception that causes askew internal representations of space. One possible explanation is hemispheric lateralisation, which could underlie such biases in spatial representation by weighting spatial signals differently across hemispheres or body sides. Supporting the idea that individual variation in spatial processing can be meaningful rather than mere noise, a recent study on *Drosophila* found individual differences in balancing stability and flexibility of spatial representations, suggesting they may arise from past experience or reflect an evolutionary strategy in which different representations confer different advantages [45]. In our study, however, past experience in video gaming did not correlate with the size of the asymmetric error component. Whether askew spatial representations might confer an evolutionary advantage in humans remains an open question for future research.

Irrespective of the perceptual system in which PI errors manifest themselves, they can arise either because walked path segments and their configuration are incorrectly coded or because the calculated target trajectory is not executed correctly [3]. In fact, we provide evidence that in our desktop VR experiment, errors arise primarily in encoding the walked path into an internal representation. Asymmetries in our experiment are unlikely to originate from execution errors, considering that there is no difference in motor control between left and right-hand trials in our study, except for the marginal difference of pushing either the left or right arrow key for turning. Consistently, we found no correlation between handedness and the size of the asymmetric error component in our study. Recent work by Chrastil and Warren provides evidence for the error source in angular PI in their experiment to be the execution of the return, because errors in their angle reproduction task depended on the magnitude of the response angle rather than the encoded angle [3]. However, Chrastil and Warren differentiated between encoding and execution errors based on responses given by left or right turns aligning parallel or opposite to a previously walked path. Our study suggests that the discrepancies between parallel and opposite responses might also originate from asymmetries between these left and right turns of their participants.

At the representational level, different weighting of outbound legs has been proposed in PI models [4]. In line with this, we propose that turns to one side might be represented with a different gain factor than turns to the other side in PI processing. Such a rotation gain factor, although not independent for each rotation direction, was recently also incorporated in a model by Castegnaro et al., who successfully used it to differentiate two subgroups of patients with mild cognitive impairments [14]. Possibly, individual asymmetries in sensitivity of motion detection might cause different angular encoding gains depending on rotation direction. At the neural level, this effect might arise from sensitivity differences of neural populations similar to asymmetric angular head velocity cells that have been identified in rats [46] and flies [47]; a hypothesis that remains to be tested in humans. Such coding might be supported by recently discovered neural populations that encode head-direction signals in humans [48, 49].

Beyond individual mechanisms, environmental features also play a role. Since our experiment focuses on visual PI, we provide only those environmental features necessary to create optic flow and to make the task sensible. If salient landmarks had been included, we would expect them to largely suppress the effects of symmetric and asymmetric PI error components[6, 7]. However, previous research has found evidence suggesting that PI biases may persist in the presence of landmark cues if landmarks are highly unreliable [15]. When experimental manipulations set PI in conflict with landmarks, as done for example by Zhao and Warren [7], or Chen et al. [6], asymmetric angle estimations cause cue conflicts of different degrees to arise for left and right trials. In environments where landmarks are unreliable, such as in deserts or dense forests, PI biases may significantly impact orientation abilities and potentially increase the risk of getting lost.

In summary, the presence of persistent directional biases may seem surprising, yet they could confer evolutionary advantages. Speculative explanations for why performance in left-hand versus right-hand triangles is not mirrored span several domains. Sensorimotor factors include visual field asymmetries, eye dominance, and postural misalignments. Cognitive factors may involve attentional asymmetries, mental rotation abilities, differential gains applied to left and right rotations by neural head-velocity populations, or hemispheric lateralisation. Finally, strategic factors, such as participants’ preferred reference frames (egocentric vs allocentric) and their alignment with task demands, may also play a role. Importantly, similar PI biases may arise from distinct underlying mechanisms. Although this study does not directly address a shared mechanism underlying asymmetric error components across active-walking and joystick paradigms, this remains an intriguing question for future work. Further research would benefit from systematically probing additional influences on PI side preferences beyond rotational perception and motor execution, for instance through manipulations of individual feedback.

### 3.4 Distance errors might arise from direction errors

We extended the simple angle estimation tasks used by Jetzschke et al. to a more complex triangle completion paradigm, revealing striking effects of rotational PI asymmetries also on distance estimation and the performance of a more complex spatial homing task [2]. Direction and distance components are correlated, or show a correlation trend, for symmetric (*p <* 0.001, *r*^2^ = 42%) and asymmetric (*p* = 0.073, *r*^2^ = 12%) error components. Unlike Harootonian et al., we also found a correlation between direction and distance errors (no symmetric or asymmetric calculations included) (supplement Fig. S5) [4]. Considering that a perceptual rotation gain in the encoding of turns (rather than in the execution, as elaborated above; see section ‘Origins and implications of path integration biases’) might cause skewed internal representations of left and right angles, this also impacts distance representation and execution because angles and distances are naturally coupled in Euclidean space (Fig. 9).

**Figure 9.**
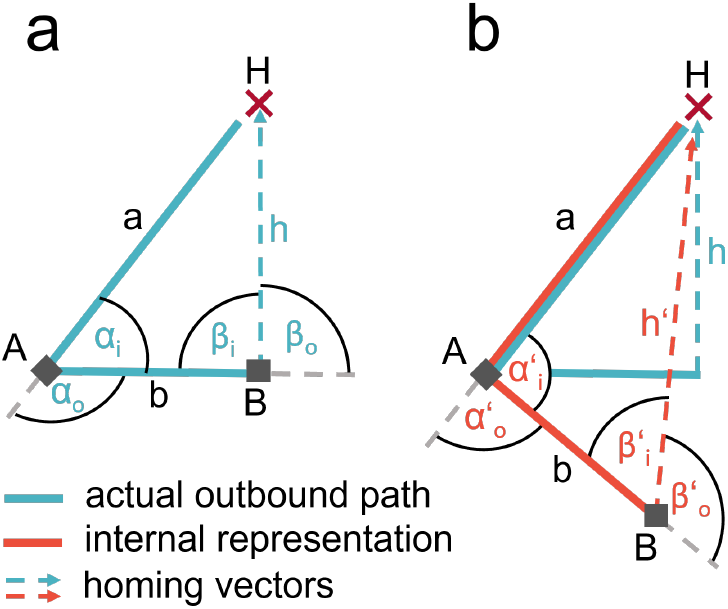
An incorrect internal representation (b, red) of actually walked angles (a, blue) affects also walked distances. Assuming an ideal internal representation of the outbound path (blue) it is geometrically unambiguously possible to infer the necessary angle to return home perfectly. A directional overshoot of the correct homing angle 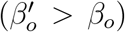 can be translated into an internal representation where the turned angle between the outbound legs (*α*_*o*_) was underestimated 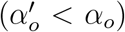. Such internal representation results in a longer executed homing distance.

Let us consider an example participant with positive symmetric direction error component, meaning the person overshoots the correct homing angle 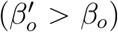 (Fig. 9). As the participant is guided along two legs of the triangle with an angle in between, it is geometrically unambiguously possible to infer the necessary angle to return home perfectly (by the law of cosine). Assuming correct calculation of the homing vector (*h*), we translate a directional overshoot of the correct homing angle (*β*_*o*_) into an internally represented underestimation of the turned angle between the outbound legs 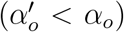. Interestingly, this underestimation of the actually turned angle also results in a longer homing vector in the internal representation (*h*^*′*^). This relation of angles and distances in Euclidean space is reflected in our experiment in correlated direction and distance errors (supplement Fig. S5) and their respective symmetric and asymmetric error components (Fig. 8). For the triangle shape tested in this study, a positive symmetric direction error component infers a positive symmetric distance error component, and vice versa for negative error component sizes. The same applies to asymmetric direction error components causing asymmetric distance error components. However, both the strength and slope of this correlation likely depend on the specific properties of the navigated path, including its shape and number of segments. Future studies can further discover the exact relation between directional and distance error components by quantifying asymmetrical error components in more triangle shapes than this study.

### 3.5 Conclusion

Our study reveals critical insights into human path integration, particularly emphasising its nuanced nature of inter-individual differences. By carefully analysing left and right trials separately in a triangle completion task, we identified individual side biases in rotation and translation that persist over time and across different geometries. The distinction between population trends and individual patterns in this study serves two purposes. Firstly, the disclosed path integration biases contribute to our understanding of human navigation and why some people might get lost in challenging environments. Secondly, our findings suggest that future studies should account for side-dependent individual biases and be mindful about variations along other experimental dimensions. To mitigate these effects, future studies should either limit the number of experimental dimensions, or collect sufficient data across all relevant dimensions. In the context of path integration studies, this could entail conducting trials for only one side and acknowledging that responses may contain some degree of bias, or collecting sufficient data across all sides and paths to contribute to a more nuanced understanding of human navigation.

## 4 Methods

### 4.1 Experimental design

This study explores individual PI error components in three parts: one part to assess time persistence in a VR triangle completion task, one part to assess persistence across path shapes therein, and one part to assess generalisability to other tasks by re-analysing data from published human navigation studies.

### 4.2 Re-analysis of data from published studies

To analyse the existence of individual symmetric and asymmetric error components in human navigation literature, we collected data of 11 published studies whose experimental design included some form of left and right-hand trials.

We performed a systematic literature search, following the basic procedure reported in the PRISMA 2020 statement to find published studies in English irrespective of publication year [42] (see supplement Fig. S1 for flow diagram). A search on the Web of Science Core Collection looking for the word combination of ‘angle’ and ‘path integration’ or the combination of ‘homing’, ‘navigation’, and ‘human’ in the title, abstract, or author keywords, conducted on 30th August 2022, yielded 451 publications. By screening titles and abstracts for general relevance 39 publications remained. We then did full-text searches to include only studies with original data and studies with individual left and right-hand trials. Screening was conducted by JS and did not include any automation tools. This procedure left us with 25 publications, applicable for our planned re-analysis, with four of them manually added by us from memory [7, 11, 33, 41]. To focus our re-analysis on the most influential studies, the 25 publications were then sorted according to their average citation count on Google Scholar, Web of Science, and Researchgate. From publications with most to fewest average citations, we then iteratively checked for data availability online, or asked the corresponding author via email, and included the dataset in our re-analysis. If no data was available, we added the publication with next most citations to our list. In total, only one single dataset was openly accessible and well documented online [4]. Ten datasets (inlcuding one former study from our own lab) were available upon request, although for the most part sparsely documented and hence only usable for re-analysis after personal clarifications by the authors. Six, relatively old (published 1991-2007), datasets were unavailable upon request with authors of one publication not responding at all. As a result we collected 11 datasets from Wiener and Mallot (2006)[1], Wiener et al. (2011)[39], Zhao and Warren (2015)[7], Jetzschke et al. (2016)[2], Petrini et al. (2016)[33], Chen et al. (2017)[6], He and McNamara (2018)[40], Lakshminarasimhan et al. (2018)[41], Sjolund et al. (2018)[11], Harootonian et al. (2020)[4], and Chrastil and Warren (2021)[3].

For the collected studies we analysed symmetric and asymmetric PI error components on an individual level using directional errors, as we calculate them in this study (see ‘Error component calculations’). Since we re-analysed PI biases directly from raw data, risks of reporting bias are inapplicable to our re-analysis. Note that we analysed each study’s data independently rather than conducting a meta-analysis. Since our goal was to examine individual PI errors for each study separately, rather than calculate a combined effect size across studies, a formal assessment of collective effect size was not necessary for our research objective. If studies included multiple conditions with the same participants, we calculated PI error components for each condition, and averaged them afterwards, resulting in one measure per participant per study. Although averaging across conditions cancels out the reported error components to some degree, we focus on the general collective evidence on individual side tendencies in PI for the sake of clarity. Details on the analysis of every single publications are given in the supplementary material (see supplement S1).

### 4.3 Experimental procedure

In addition to the re-analysis of data from earlier studies we conducted an experiment dedicated to analyse error components in PI at the individual level. *N* = 27 healthy participants (aged 27.5±8.8 (mean ± sd) years, 18 self-identified as female, 9 self-identified as male) took part in the experiment. Participants were recruited from the department’s online participant portal, which is accessible to the general public.. Participants were required to be at least 18 years old (the legal age of adulthood in Germany) and to have normal or corrected-to-normal vision and hearing. No additional specific inclusion or exclusion criteria were applied. Participants were informed about the general procedure of the experiment but not about its specific purpose or the nature of the different conditions. All participants gave their written informed consent to participate and received monetary compensation (10 Euros per hour). The experiments were approved by the Bielefeld University Ethics Committee under the reference number EUB 2019-174 and conducted in accordance with the guidelines of the Deutsche Gesellschaft für Psychologie e.V. (DGPs), which correspond to the guidelines of the American Psychological Association (APA). All data collected from the participants and during the experiment was handled according to the General Data Protection Regulation (GDPR) of the European Union. Participants were invited to the lab three times. On the first day, they conducted the VR triangle completion task with triangle shape A (see ‘Virtual reality triangle completion task’ for details). On the second day (min= 5, mean= 21.7 days later), they conducted the same task with triangle shape B. On the third day (min= 21, mean= 35.7 days later), they conducted the VR task with triangle shape A again. Eight participants were not tested with triangle shape B. We focus analyses on error component persistence across triangle shapes on *N* = 19 participants (aged 27.6 ± 10.1 (mean ± sd) years, 13 self-identified as female, 6 self-identified as male).

The VR experiments of each day were carried out over three sessions (1 training + 2 test sessions). The experiment was provided in the form of a stand-alone Windows application, created using the ‘Virtual Navigation Toolbox’ for the Unity3D game engine [50] and was executed on a desktop computer. An experimenter was present to instruct and assist participants according to a standardised protocol during the initial six training trials. After training, participants completed the remaining test sessions without further supervision. However, participants were instructed to take regular breaks in between experimental sessions of at least 5-10 minutes. Sessions on average took 28.2 ± 4 minutes to complete. Each test session consisted of 15 repetitions per side (left-hand or righthand triangle) of one triangle shape (A or B) in randomised order, leading to a total of 30 trials per session and adding up to overall *n* = 30 repetitions per triangle side for each participant. The high number of repetitions allows to well estimate individual response accuracy and precision. Testing the triangle on both sides is necessary to calculate the symmetric and the asymmetric individual error components (see section ‘Error component calculations’ for details).

### 4.4 Virtual reality triangle completion task

We tested participants in a desktop VR large-scale triangle completion task designed with the ‘Virtual Navigation Toolbox’ (Fig. 1a) [50].

In our VR version of the task, the participants were first guided to their home location indicated by a stylised campfire. This initial approach always started 10 virtual meters (*vm*) behind the campfire (Fig. 4). The avatar was controlled from a first-person view using the up/down and the left/right arrow keys, respectively, for translation and rotation. The campfire then disappeared and an arrow at the bottom of the screen guided the participant to collect firewood logs at two distinct locations, while being able to move freely. This was termed the outbound phase of the trial. To ensure an overall straight path along the legs of the outbound path, the participant was confronted with a time limit (40 *s*) to collect all the firewood. The remaining time was always visible in the form of a timeout bar at the top of the screen. After collecting the second firewood log, participants were again presented with a 40 *s* timer, and had to find back to the campfire location (i.e. the goal location), entering their response via button press. Neither the firewood logs nor the campfire were visible during the return, to prevent their use as spatial cues. After the participant had indicated reaching the goal, the next trial was started by teleporting the avatar to the start location 10 *vm* behind the campfire, while at the same time the environment for the new trial was regenerated. No feedback on performance was given. During each trial, the position and rotation of the player’s avatar in the virtual environment were recorded for every rendered frame. Identical grass tufts on the ground were randomly arranged for each trial to enable visual estimation of self-motion by optic flow. A sun-like light source provided global illumination from directly above so that no directional information could be derived from shadows in the environment. The avatar moved at 2.5 *vm* per second and turned at 60° per second. Avatar and objects were scaled to make virtual meters resemble real meters.

Before the test session, participants conducted 6 training trials, for experimenters to ensure the task and controls to be well understood (one participant conducted 17 training trials until the avatar control was completely natural). Training trials were equal to test trials, except for two adaptations. First, after reaching the campfire, participants were allowed to freely explore the environment for 90 *s*. Second, when the homing phase of the trial starts at the second firewood log neither the guidance arrow at the bottom of the screen, nor the campfire disappeared, but participants were guided by the arrow to the campfire, instead of having to remember its position and navigate there on their own. This training procedure calibrates participants for the correct turning angle and should compensate for individual differences in perceiving the VR. Therefore, any remaining individual error components persistent across time or triangle shape should not be the result of artifacts caused by the technical details of the experimental design.

The study was conducted in two parts, one part testing time persistence with the same triangle shape on two different days, and one part testing shape persistence with two different triangle shapes.

The first study part was conducted with two sessions in triangle A, conducted at least three weeks apart. The firewood logs that indicate the triangle corners were located at *A* = (20, 20), or *A* = (−20, 20), for left-hand and right-hand trials respectively, and *B* = (0, 20), with the campfire home location at the origin (0, 0) (Fig. 4a and b). Total path length was 68.28 *vm*. At location B, participants had to rotate 90° left or right for the correct heading-direction towards the home location.

The second study part was conducted by *N* = 19 of the same participants with one session in triangle A, as described above, and one session in a more pointed triangle B. The firewood logs were located at *A* = (0, 12.07) and *B* = (−36.21, 12.07), or *B* = (36.21, 12.07), for left-hand and right-hand trials respectively, with the campfire home location at the origin (0, 0) (Fig. 4c). Total path length was 63.05 *vm*, with the second outbound leg being three times longer than the first. At location B, participants had to rotate 161.6° left or right for the correct heading-direction towards the home location.

### 4.5 Data analysis

To analyse persistent errors in human PI at various experimental dimensions, we calculate symmetric and asymmetric individual error components based on signed direction or distance errors from triangle completion tasks (Fig. 1c and d).

To resolve differences between turning direction, participants are tested in triangles including either only left turns, termed ‘left-hand triangle’, or only right turns, termed ‘right-hand triangle’ (Fig. 1b). In a single trial, participants walk from home location *H* along the first triangle leg *a* to corner *A*, then turn through the outer angle *α*_*o*_ (the opposite of the inner angle *α*_*i*_), and walk along the second triangle leg *b* to arrive at the second corner *B* (Fig. 1b). At this point, participants are asked to freely return to *H*. The correct homing angle *β*_*o*_ (opposite of the inner triangle angle *β*_*i*_) leads to the correct homing trajectory *h*_*c*_. We observe homing trajectories *h*_*o*_ deviating from *h*_*c*_. We calculate the signed direction error as the angular deviation between the correct and the observed homing direction, based on the final response position of each trial. The direction error is signed and defined as positive for deviations to the right, and as negative for deviations to the left, from a participant’s ego perspective.

All analyses were conducted in custom Python scripts in version 3.11.9 [51].

#### 4.5.1 Error component calculations

From the observed *n* = 30 measurements of direction errors in randomly shuffled left and right-hand triangles, we calculate distributions of the symmetric and asymmetric error components at individual participant level as follows. The degree of symmetric error component (*err*_*sym*_) is calculated by:

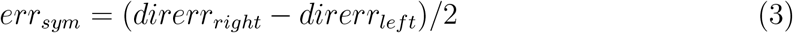

where *direrr*_*right*_ is one direction error in a right-hand triangle trial and *direrr*_*left*_ is one direction error in a left-hand triangle trial. This component is the shared degree of over- or underturning in left and right turns. Figuratively, one can illustrate the symmetric error component as half of the angle between the walked directions in a left-hand and a right-hand trial (Fig. 1c, in dark magenta).

The asymmetric error component (*err*_*asym*_) is calculated by:

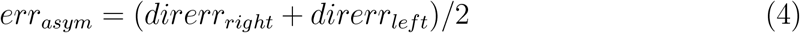

This component describes the tendency of a participant towards the left (negative sign) or right (positive sign) side. One can think of the asymmetric error component as the difference between the tendencies to over- or undershoot left or right turns. Pictorially, one can illustrate the asymmetric error component as half of the angular difference between the vector pointing into the walked direction of a right-hand trial, and the projection of the vector of a left-hand trial to the other side (Fig. 1c, in light magenta).

#### 4.5.2 Statistics

A value for the symmetric or the asymmetric error component can only be calculated from the combination of one left-hand with one right-hand direction error. As direction error measurements for left and right-hand triangles are independent, we pair each of the *n* = 30 repetitions per side with each other to eventually calculate a distribution of 900 values, each for the symmetric and for the asymmetric error component. The mean of the generated distribution is equivalent to the error component size when calculating it based on the mean direction error of all left and of all right-hand trials. We use the distributions to calculate confidence intervals (ci) and conduct two-sided bootstrapping hypothesis testing following the basic procedure described by Bittmann [52]. All reported error component sizes are based on circular calculations.

To analyse the existence of both error components at the individual level, we test the size of each participant’s error component against the null hypothesis of it being equal to zero with bootstrapped ci. We report the proportion of participants showing error component sizes that are significantly different from zero for one or both experiment days (Fig. 5c and 7c). To analyse time persistence of both error components, we plot all participants’ error component sizes for each of the two experiment days along with the degree of change between the days and a correlation between session one and session two (Fig. 5a, b, d, e, 7a, b, d, e). Based on these distributions, we compare variance across participants between the symmetric and the asymmetric error component. To analyse persistence across triangle shapes, we conduct the same analysis as for time persistence for the two different triangle shapes in our experiment, and plot error component sizes for each triangle along with the degree of difference, as well as correlations between triangle A and triangle B (Fig. 6). Cohen’s *d* effect sizes calculated by (*x* − *µ*)*/σ* are indicated where applicable. Sample sizes and test statistics are indicated where applicable throughout the Results.

#### 4.5.3 Distance biases

Besides error components in direction error, we can also calculate symmetric and asymmetric error components of distance estimation. The distance error is the deviation from the correct homing distance and defined as positive for deviations where participants walk too far and as negative for deviations where participants walk too short (Fig. 1d). The distance error is calculated based on the final location of a participant. The symmetric error component for distance describes the tendency of a participant to over- or undershoot the correct homing distance symmetrically for left and right-hand triangles. The asymmetric error component for distance describes the tendency of a participant to over- or undershoot the correct homing distance for one triangle side systematically more than for the other. When the asymmetric distance error is positive, it indicates that the person tended to walk further in right-hand triangles; a negative value reflects the opposite pattern. To check for the existence and features of symmetric and asymmetric distance error components we execute the same statistical analyses as for the direction error described above (see section ‘Statistics’).

To analyse reciprocal influences, we set the distance and direction error components into relation and quantify correlations between both measures (Fig. 8).

#### 4.5.4 Data cleaning

We excluded trials for the following four reasons. First, when participants turned into the wrong direction at homing start, namely in counter-clockwise direction for right-hand triangles, or in clockwise direction for left-hand triangles. Second, due to technical issues some trials do not possess a homing trajectory or the response of participants was given exactly at the homing start location. Third, when participants walked backwards and kept the campfire home location in sight for the whole duration of the trial. Fourth, due to a mistake in the session with triangle shape B, for some participants two trials in these sessions were guiding participants through triangle shape A. We did not exclude any outliers from analysis. On average, 2.9 trials per participant had to be excluded from analysis per experiment.

### 4.6 Power analysis

Based on response variance in one of our previous studies, that contained a very similar condition as used here [15], we performed a power analysis to estimate the number of repetitions per side needed to resolve performance differences of a given magnitude. Based on one-sample t-tests to be able to detect symmetric and asymmetric error components of 2.5° with a power of 80%, we would have needed a minimum sample size per triangle side of *n* = 25. Based on that calculation, we eventually chose to do *n* = 30 repetitions per side.

During our final analyses on circular direction errors we did not rely on the assumption of normality, and switched to bootstrapping hypothesis testing which in a post-hoc analysis with 30 repetitions per side revealed that we can, in this study, resolve differences in both error components of 2° with 85% power, and of 3° with 98% power.

### 4.7 LLM usage

We utilised GPT-4 from OpenAI and Claude 3.5 Sonnet from Anthropic during the writing procedure of this manuscript.

## 5 Acknowledgement

We want to thank all authors for their support in providing data and clarifying notes on data structures for the re-analysis of published studies. We would like to highlight the large effort that was necessary to collect data of publications in our field of human navigation research. We strongly encourage all scientists to provide their data in an openly accessible way accompanied by appropriate documentation, to facilitate data re-use and conduction of meta-analyses of different forms. We acknowledge the financial support of the Deutsche Forschungsgemeinschaft (DFG) (grant 460373158 https://gepris.dfg.de/gepris/projekt/460373158). The funders had no role in study design, data collection and analysis, decision to publish, or preparation of the manuscript. We would also like to thank the reviewers involvesd in the parallel publication process for their careful reading of our manuscript and for their insightful comments and suggestions, which helped us to improve the manuscript considerably.

## 6 Data Availability

The datasets generated and analysed during the current study are available in Bielefeld University’s PUB repository at https://doi.org/10.4119/unibi/3006526. Analysis code with accompanying instructions for execution is available on GitLab at https://gitlab.ub.uni-bielefeld.de/jonas.scherer/uncovering_persistent_biases_in_human_path_integration_by_separating_left_and_right_trials. Implementation details of the Unity3D project and an executable Unity project of this study’s task are available on GitLab at https://gitlab.ub.uni-bielefeld.de/virtual_navigation_tools/unity_vnt_showcase_triangle_completion.

## 7 Contributions

Conceptualisation: J.S., M.M.M., O.J.N.B., and N.B.; Methodology: J.S., and M.M.M.; Software: M.M.M.; Investigation: A.K.; Data curation: J.S., and A.K.; Visualisation: J.S.; Formal Analysis: J.S.; Validation: A.K.; Writing-original draft: J.S.; Writing-review & editing: J.S., M.M.M., A.K., M.E., O.J.N.B., and N.B.; Funding acquisition, project administration, resources, and supervision: M.E., O.J.N.B., and N.B..

