## Supplementary Material for "Uncovering persistent biases in human path integration by separating left and right trials"

Jonas Scherer [1,*], Martin M. Müller [1], Anabel Kroehnert [2], Martin Egelhaaf [1], Olivier J. N. Bertrand [1,] and Norbert Boeddeker [2,]
[1]Department of Neurobiology, Bielefeld University, Germany
[2]Department of Cognitive Neuroscience, Bielefeld University, Germany

#### 1 S1: Re-analysis of data from published studies

To analyse the existence of individual symmetric and asymmetric error components in influential human navigation literature, we used the basic procedure reported in the PRISMA 2020 statement [1] to collect data of 11 published studies whose experimental design included some form of left and right-hand trials. Here we present the PRISMA flow diagram for the identification of studies (Fig. S1) and describe for each study how exactly we re-analysed the data to calculate error component sizes at inidividual level. Studies are listed alphabetically. Details about inclusion and exclusion criteria are given in the main text (section "Re-analysis of data from published studies")

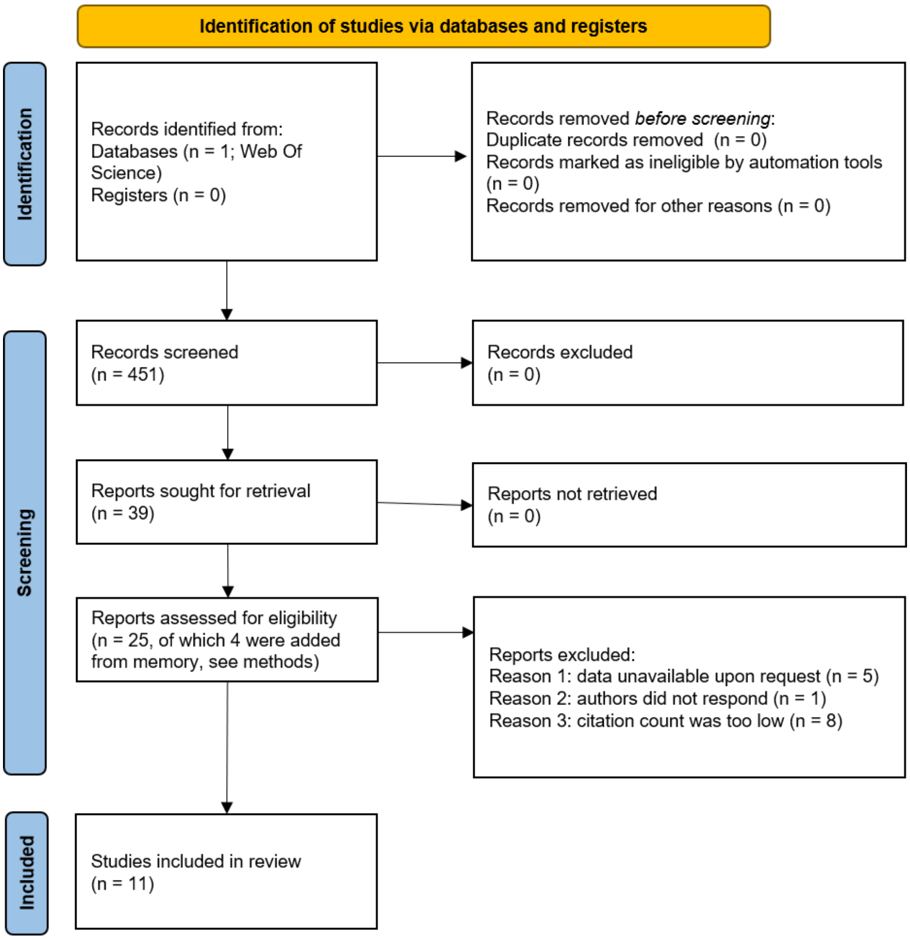


Figure S1: Flow diagram of identification of studies via databases and registers from the PRISMA 2020 statement [1]. To focus our re-analysis on the most influential studies, the 25 eligible publications were sorted according to their average citation count on Google Scholar, Web of Science, and Researchgate. From publications with most to fewest average citations, we iteratively checked for data availability online, or asked the corresponding author via email, and included the dataset in our re-analysis. If no data was available, we added the publication with next most citations to our list.

#### 1.1 Chen et al., 2017

Chen et al. (2017) [2] conducted triangle completion experiments in virtual reality presented by a head-mounted display. Participants performed the task by active walking. For our re-analysis we focus on the data of the "self-motion" condition in Experiment 1a because it does not contain visual landmarks and does not manipulate the reliability of self-motion cues. We calculate the direction errors separately for left and right-hand triangles, defined by the configuration of the posts, from the deviation of the correct homing vector and the response vector. We define direction errors as positive for deviations to the right and as negative for deviations to the left from the participant's ego perspective. Following the calculations outlined in our main manuscript we calculate the symmetric and asymmetric error component separately for each participant and separately for the "rich" and ”poor” environment which, however, had the same properties for the ”self-motion” trials nonetheless (see ”Error component calculations” in ”Methods”). For plotting, we averaged individual’s error component sizes across the ”rich” and ”poor” condition and indicate each of $n = 18$ participant’s mean as one datapoint (Fig. 2).

#### 1.2 Chrastil and Warren, 2021

Chrastil and Warren (2021) [3] conducted triangle completion experiments and angle reproduction tasks in virtual reality presented by a head-mounted display. Participants performed the tasks by active walking. For our re-analysis we focus on the data of the triangle completion tasks in the ”hallway” and ”open field” conditions. Other than in the ”open field” condition, in the ”hallway” condition participants were surrounded by elongated or circular hedges. Both conditions contained trials in six different triangle path configurations. We define direction errors as positive for deviations to the right and as negative for deviations to the left from the participant’s ego perspective. Following the calculations outlined in our main manuscript we calculate the symmetric and asymmetric error component separately for each participant, separately for the ”hallway” and ”open field” condition, and separately for each triangle shape therein (see ”Error component calculations” in ”Methods”). For plotting, we average individual’s error component sizes across the ”hallway” and ”open field” condition and across all triangle shapes and indicate each of $n = 13$ partcipant’s mean as one datapoint (Fig. 2).

#### 1.3 Harootonian et al., 2020

Harootonian et al. (2020) [4] conducted triangle completion experiments in virtual reality presented by a head-mounted display. Participants performed the tasks by actively walking on an omnidirectional treadmill. For our re-analysis we consider all conditions in experiment 1, in which the authors tested particpants in seven different triangle path configurations, and all conditions in experiment 2, that contained triangles with five different spatial scales. We define direction errors as positive for deviations to the right and as negative for deviations to the left from the participant’s ego perspective. Following the calculations outlined in our main manuscript we calculate the symmetric and asymmetric error component separately for each participant and for each experiment (see ”Error component calculations” in ”Methods”). Because of the low number of repetitions per triangle shape and size (n=4), different triangle shapes and sizes were pooled for calculations right away. For plotting, we average individual’s error component sizes across both experiments and indicate each of $n = 39$ partcipant’s mean as one datapoint (Fig. 2).

#### 1.4 He and McNamara, 2018

He and McNamara (2018) [5] conducted triangle completion experiments in virtual reality presented by a head-mounted display. Participants performed the tasks by active walking. For our re-analysis we considered both, the "configural" and the "continuous" condition. We calculate the direction errors separately for left and right-hand triangles, inferred by the configuration of the posts in each trial, from the deviation of the correct homing vector and the response vector. We define direction errors as positive for deviations to the right and as negative for deviations to the left from the participant's ego perspective. Following the calculations outlined in our main manuscript we calculate the symmetric and asymmetric error component separately for each participant each of which were tested either in the "configural" or the "continuous" condition (see "Error component calculations" in "Methods"). We excluded participant 4 of the "continuous" condition because the response angle is exactly the same for the most part and errors result to be unrealistically high for that participant. For plotting, we averaged individual's error component sizes and indicate each of $n = 24$ participant's mean as one datapoint (Fig. 2).

#### 1.5 Jetzschke et al., 2016

Jetzschke et al. (2016) [6] conducted active and passive angle estimation experiments. Participants actively walked blindfolded or in a dark room while being guided or instructed to turn through angles of different degrees. For our re-analysis we consider both of these experiments. We base calculations of the symmetric and asymmetric error component on actively turned angles (experiment 1) or points of subjective equality (experiment 2) for left and right turns and use the equations outlined in our main manuscript (see "Error component calculations" in "Methods"). For plotting, we average individual's error component sizes and indicate each of $n = 7$ participant's mean as one datapoint (Fig. 2).

#### 1.6 Lakshminarasimhan et al., 2018

Lakshminarasimhan et al. (2018) [7] conducted optic flow based path integration experiments in virtual reality. Stimuli were presented on screen and participants controlled their walking via a joystick with two degrees of freedom. For our re-analysis we consider all participants in all reported conditions. Conditional variations include but are not limited to number of repetitions, linear and angular speed, panorama, optic flow density, and spatial scale (see "Method Details - Behavioural Task" for details [7]). We calculate the direction errors separately for left and right trials, inferred by the target being located at the left or right side of the participant from the deviation of the correct homing angle and the response angle. We define direction errors as positive for deviations to the right and as negative for deviations to the left from the participant's ego perspective. Following the calculations outlined in our main manuscript we calculate the symmetric and asymmetric error component separately for each participant (see "Error component calculations" in "Methods"). For plotting, we averaged individual's error component sizes and indicate each of $n = 20$ participant's mean as one datapoint (Fig. 2).

#### 1.7 Petrini et al., 2016

Petrini et al. (2016) [8] conducted a reproduction task for paths with two legs in virtual reality presented by a head-mounted display. Participants performed the tasks by active walking. For our re-analysis we focus on the data of the "self-motion only" condition because it does not contain visual landmarks. We calculate direction errors, separately for trials with left and right turns, as the deviation of the correct turning angle between the two walked legs, and the response turning angle between legs. We define direction errors as positive for deviations to the right and as negative for deviations to the left from the participant's ego perspective.

Because we were provided with only the raw trajectory data we calculated response turning angles between legs as follows. First, we find the turning point by taking the percentage where theoretically the perfect turn would be (length of leg 1 divided by length of leg 2: $2.7/(2.7+2) \approx 57\%$ and take a window of 50% around it. In this we smooth the angular velocity with a running average of size three and find its local maximum that we define as the turning point. Around the turning location we define an area of $\pm 20\,cm$ in space as neutral area that we do not take into consideration for further analyses. We also do not take into consideration the first 2% and the last 2% of distance covered. For the two legs before and after the turn we rotate datapoints by their indicated rough walking direction onto the x-axis, fit linear regressions to estimate the real walking direction, and rotate the points back. The angle between the resulting paths is the response turning angle. As turning points are only defined by the overall path shape, we excluded $n = 5$ trials because turning points were not clearly detectable:

Following the calculations outlined in our main manuscript we calculate the symmetric and asymmetric error component separately for each participant (see "Error component calculations" in "Methods"). For plotting, we averaged individual's error component sizes and indicate each of $n = 33$ participant's mean as one datapoint (Fig. 2).

#### 1.8 Sjolund et al., 2018

Sjolund et al. (2018) [9] conducted triangle completion experiments in virtual reality presented by a head-mounted display. Participants performed the task by active walking. For our re-analysis we focus on the data of the "path integration only" condition in

Experiment 1 because it does not contain visual landmarks. We calculate the direction errors separately for left and right-hand triangles, defined by the configuration of the first and second triangle post, from the deviation of the correct homing vector and the response vector. We define direction errors as positive for deviations to the right and as negative for deviations to the left from the participant's ego perspective. Following the calculations outlined in our main manuscript we calculate the symmetric and asymmetric error component separately for each participant (see "Error component calculations" in "Methods"). Because the target location in the experiment was randomly selected $n = 3$ participants were only tested on one side. The respective participants are indicated by nan entries in the accompanying dataset. For plotting, we averaged individual's error component sizes and indicate each of the remaining $n = 48$ participant's mean as one datapoint (Fig. 2).

#### 1.9 Wiener and Mallot, 2006

Wiener and Mallot (2006) [10] conducted optic flow based homing experiments in virtual reality in which participants had to point to the origin of paths with different numbers of segments. Stimuli were presented on screen and participants controlled their walking via a joystick. For our re-analysis we consider all conditions with six different overall turning angles and four different segment counts. We calculate the direction errors separately for left and right-hand triangles from the deviation of the correct homing vector and the response vector. We define direction errors as positive for deviations to the right and as negative for deviations to the left from the participant's ego perspective. Following the calculations outlined in our main manuscript we calculate the symmetric and asymmetric error component separately for each participant (see "Error component calculations" in "Methods"). For plotting, we averaged individual's error component sizes across all walked paths and indicate each of $n = 19$ participant's mean as one datapoint (Fig. 2).

#### 1.10 Wiener et al., 2011

Wiener et al. (2011) [11] conducted triangle completion experiments with blindfolded, actively walking participants. For our re-analysis we consider both, the "configural" and the "continuous" condition. In the dataset that was sent to us there was an additional "uninstructed" condition that was neither used in the original publication nore in our re-analysis. We define direction errors as positive for deviations to the right and as negative for deviations to the left from the participant's ego perspective. Following the calculations outlined in our main manuscript we calculate the symmetric and asymmetric error component separately for each participant and for each condition "configural" or "continuous" (see "Error component calculations" in "Methods"). For plotting, we averaged individual's error component sizes across all walked paths and indicate each of $n = 15$

participant's mean as one datapoint (Fig. 2).

#### 1.11 Zhao and Warren, 2015

Zhao and Warren (2015) [12] conducted triangle completion experiments in virtual reality presented by a head-mounted display. Participants performed the task by active walking. For our re-analysis we focus on the data of the "path-integration-alone" condition because it does not contain visual landmarks and does not manipulate the reliability of self-motion cues, and therein only on the "distal landmarks" condition because only six participants were tested in the "local landmark" condition. We calculate the direction errors separately for left and right-hand triangles, defined by the configuration of the posts, from the deviation of the correct homing vector and the response vector. We define direction errors as positive for deviations to the right and as negative for deviations to the left from the participant's ego perspective. Following the calculations outlined in our main manuscript we calculate the symmetric and asymmetric error component separately for each participant (see "Error component calculations" in "Methods"). For plotting, we averaged individual's error component sizes across the "rich" and "poor" condition and indicate each of $n = 12$ participant's mean as one datapoint (Fig. 2).

### 2 S2: Initial Heading

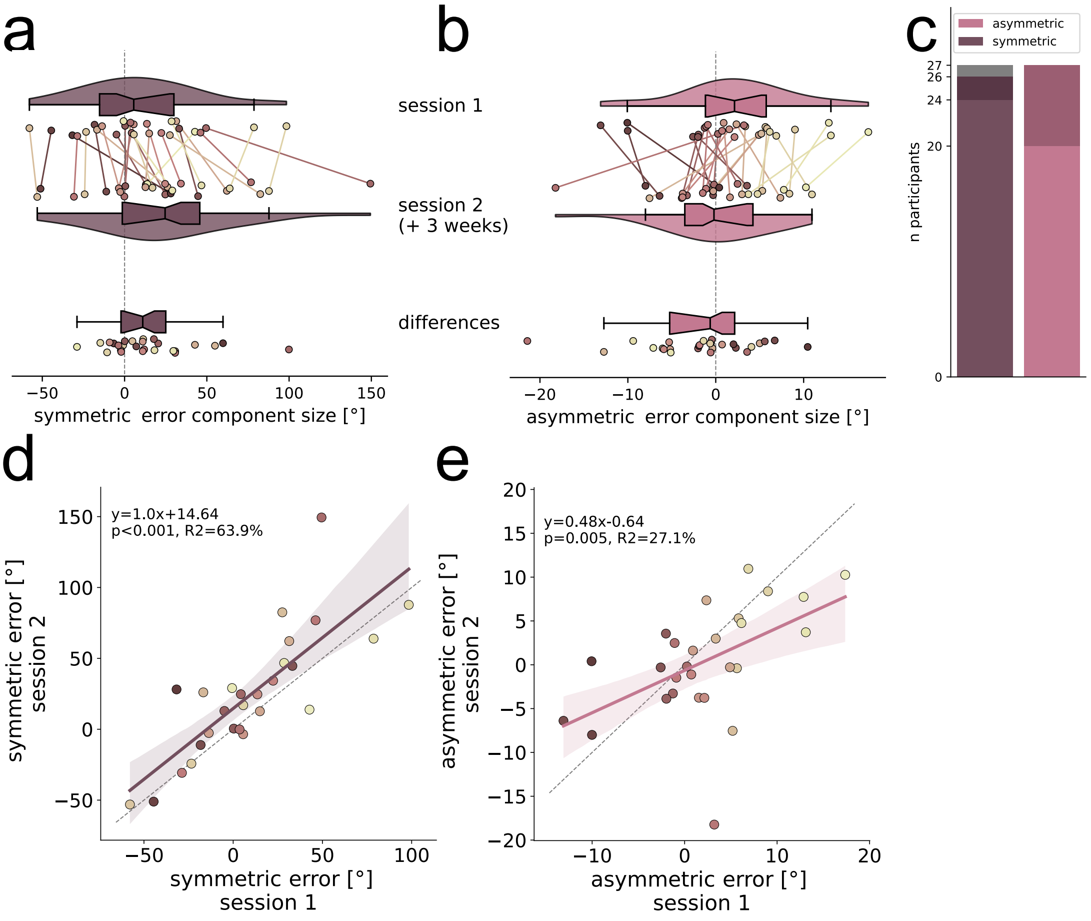


Figure S2: Symmetric and asymmetric direction error component sizes in session one and session two calculated from the initial heading direction after the first $2\,vm$ of the homing trajectory. (a) Symmetric and (b) asymmetric error component sizes, reported in °, are calculated per individual participant ($N = 27$) for each of two sessions that were conducted at least three weeks apart and are indicated by different coloured dots. Lines of the same colour connect data points of individuals across sessions and indicate the amount of change. Dot colours are sorted from dark to light, corresponding from low to high asymmetric direction error component size, and are consistent across all figures. Violins depict kernel density estimations and contain boxplots with notches indicating 95% confidence intervals. Additionally, we report the amount of difference between sessions. (c) Number of participants with significant (bootstrapped 95% confidence intervals around mean) error components in one (dark colours) or both (light colours) sessions (see section "Error component calculations" in "Methods" for details). 26/27 participants show significant error components in at least one of two sessions. (d) Symmetric and (e) asymmetric error components are persistent across sessions. Linear regressions with 95% confidence intervals are plotted in magenta (symmetric) or light magenta (asymmetric) and described by the line equation, coefficient of determination $r^2$, and Pearson correlation $p$-value.

### 3 S3: Subjective gaming experience and handedness

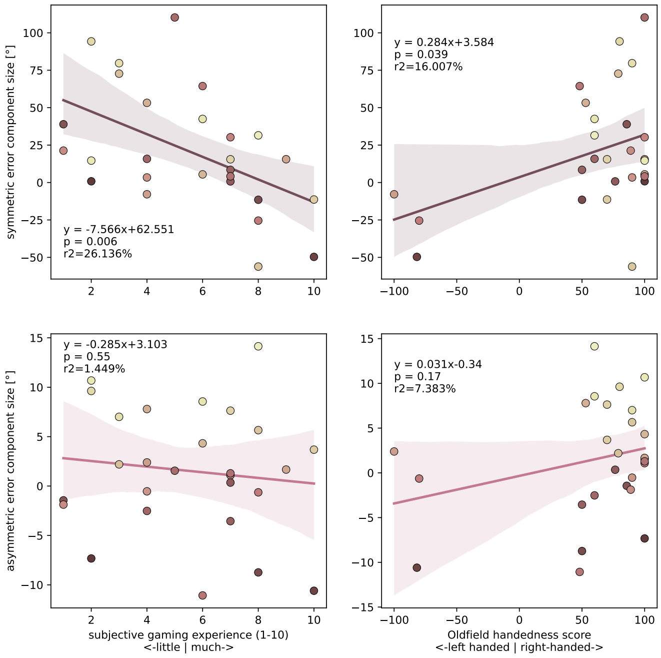


Figure S3: Correlations of symmetric and asymmetric direction error component sizes in triangle A with subjective prior experience with computer games and with participants' handedness. Symmetric and asymmetric error component sizes, reported in °, are averaged over two days per individual participant ($N = 27$) and indicated by dots. Gaming experience was assessed subjectively with a questionnaire ("How much experience do you have with pc gaming?"). To assess handedness the Edinburgh Handedness Inventory by Oldfield was used [13]. Dot colours are sorted from dark to light, corresponding from low to high asymmetric direction error component size, and are consistent across all figures. Linear regressions with 95% confidence intervals are plotted in magenta (symmetric) or light magenta (asymmetric) and described by the line equation, coefficient of determination $r^2$, and Pearson correlation $p$-value.

### 4 S4: Triangle B distance error components

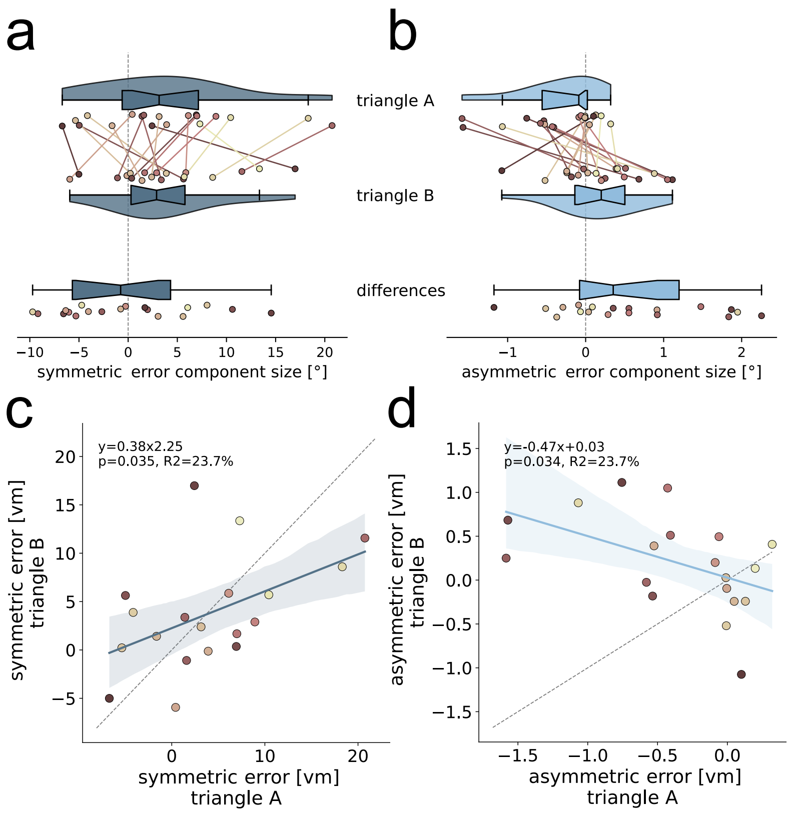


Figure S4: Symmetric and asymmetric distance error component sizes in triangle A and triangle B. (a) Symmetric and (b) asymmetric error component sizes, reported in $vm$, are calculated per individual participant ($N = 19$) for each of two triangle path configurations and are indicated by different coloured dots. Violins of triangle A averages across both experimental sessions conducted at least three weeks apart. Data for triangle B was collected in only one session. Only participants who were tested in both triangles are shown. Lines of the same colour connect data points of individuals across sessions and indicate the amount of change. Dot colours are sorted from dark to light, corresponding from low to high asymmetric direction error component size, and are consistent across all figures. Violins depict kernel density estimations and contain boxplots with notches indicating 95% confidence intervals. Additionally, we report the amount of difference between sessions. (c) Symmetric, but not (d) asymmetric error component, is persistent across triangle path configurations. Linear regressions with 95% confidence intervals are plotted in dark blue (symmetric) or light blue (asymmetric) and described by the line equation, coefficient of determination $r^2$, and Pearson correlation $p$-value.

### 5 S5: Direction and distance error correlation

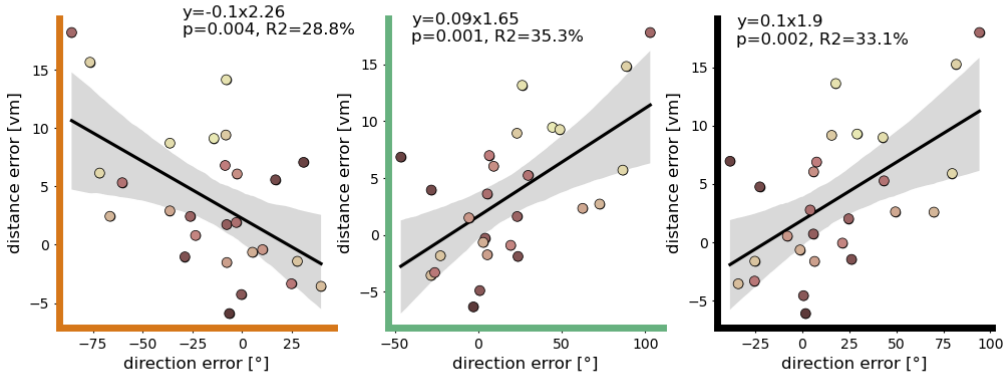


Figure S5: Direction and distance errors of each individual participant ($N = 27$) are correlated for left-hand, and right-hand trials, and for pooled sides with one side mirrored to the other. Scattered datapoints show mean direction errors, in ° on the x-axis, and distance errors, in $vm$ on the y-axis, of single participants in left-hand trials (left, orange), right-hand trials (middle, green), and pooled sides (right, black). For pooled datapoints, direction errors of left-hand trials were inverted. Dot colours are sorted from dark to light, corresponding from low to high asymmetric direction error component size, and are consistent across all figures. Linear regressions with 95% confidence intervals are plotted in dark blue (symmetric) or light blue (asymmetric) and described by the line equation, coefficient of determination $r^2$, and Pearson correlation $p$-value.
